# PhyDOSE: Design of Follow-up Single-cell Sequencing Experiments of Tumors

**DOI:** 10.1101/2020.03.30.016410

**Authors:** Leah Weber, Nuraini Aguse, Nicholas Chia, Mohammed El-Kebir

## Abstract

The combination of bulk and single-cell DNA sequencing data of the same tumor enables the inference of high-fidelity phylogenies that form the input to many important downstream analyses in cancer genomics. While many studies simultaneously perform bulk and single-cell sequencing, some studies have analyzed initial bulk data to identify which mutations to target in a follow-up single-cell sequencing experiment, thereby decreasing cost. Bulk data provide an additional untapped source of valuable information, composed of candidate phylogenies and associated clonal prevalence. Here, we introduce PhyDOSE, a method that uses this information to strategically optimize the design of follow-up single cell experiments. Underpinning our method is the observation that only a small number of clones uniquely distinguish one candidate tree from all other trees. We incorporate distinguishing features into a probabilistic model that infers the number of cells to sequence so as to confidently reconstruct the phylogeny of the tumor. We validate PhyDOSE using simulations and a retrospective analysis of a leukemia patient, concluding that PhyDOSE’s computed number of cells resolves tree ambiguity even in the presence of typical single-cell sequencing errors. We also conduct a retrospective analysis on an acute myeloid leukemia cohort, demonstrating the potential to achieve similar results with a significant reduction in the number of cells sequenced. In a prospective analysis, we demonstrate that only a small number of cells suffice to disambiguate the solution space of trees in a recent lung cancer cohort. In summary, PhyDOSE proposes cost-efficient single-cell sequencing experiments that yield high-fidelity phylogenies, which will improve downstream analyses aimed at deepening our understanding of cancer biology.

**Author summary:** Cancer development in a patient can be explained using a phylogeny — a tree that describes the evolutionary history of a tumor and has therapeutic implications. A tumor phylogeny is constructed from sequencing data, commonly obtained using either bulk or single-cell DNA sequencing technology. The accuracy of tumor phylogeny inference increases when both types of data are used, but single-cell sequencing may become prohibitively costly with increasing number of cells. Here, we propose a method that uses bulk sequencing data to guide the design of a follow-up single-cell sequencing experiment. Our results suggest that PhyDOSE provides a significant decrease in the number of cells to sequence compared to the number of cells sequenced in existing studies. The ability to make informed decisions based on prior data can help reduce the cost of follow-up single cell sequencing experiments of tumors, improving accuracy of tumor phylogeny inference and ultimately getting us closer to understanding and treating cancer.

## Introduction

Tumorigenesis follows an evolutionary process during which cells gain and accumulate somatic mutations that lead to cancer [1]. The most natural expression of an evolutionary process is a *phylogeny* — a tree that describes the order and branching points of events in the history of a cellular population. Tumor phylogenies are critical to understanding and ultimately treating cancer, with recent studies using tumor phylogenies to identify mutations that drive cancer progression [2,3], assess the interplay between the immune system and the clonal architecture of a tumor [4,5], and identify common evolutionary patterns in tumorigenesis and metastasis [6,7]. These downstream analyses critically rely on accurate phylogenies that are inferred from sequencing data of a tumor.

The majority of current cancer genomics data consist of pairs of matched normal and tumor samples that have undergone bulk DNA sequencing. Bulk data is composed of sequences from cells with distinct genomes. More specifically, we observe frequencies **f** = [*f_i_*] for the set of somatic mutations in the tumor (Fig 1a). Many deconvolution methods have been proposed for tumor phylogeny inference from such data [8–13], typically inferring a set 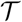 of equally plausible trees (Fig 1b). These approaches are unsatisfactory, as candidate trees with different topologies may alter conclusions in downstream analyses. Single-cell sequencing (SCS), as opposed to bulk sequencing, enables us to observe specific clones present within the tumor. These clones correspond to the leaves of the true phylogeny, allowing phylogeny inference methods to reconstruct the tree itself once we observe all clones in the tumor [14–17]. However, the elevated error rates of SCS, as well as its high cost [18], make it prohibitive as a standalone method for phylogeny inference. As such, hybrid methods have been recently proposed to infer high-fidelity phylogenies from combined bulk and SCS data obtained from the same tumor [19,20].

**Fig 1.**
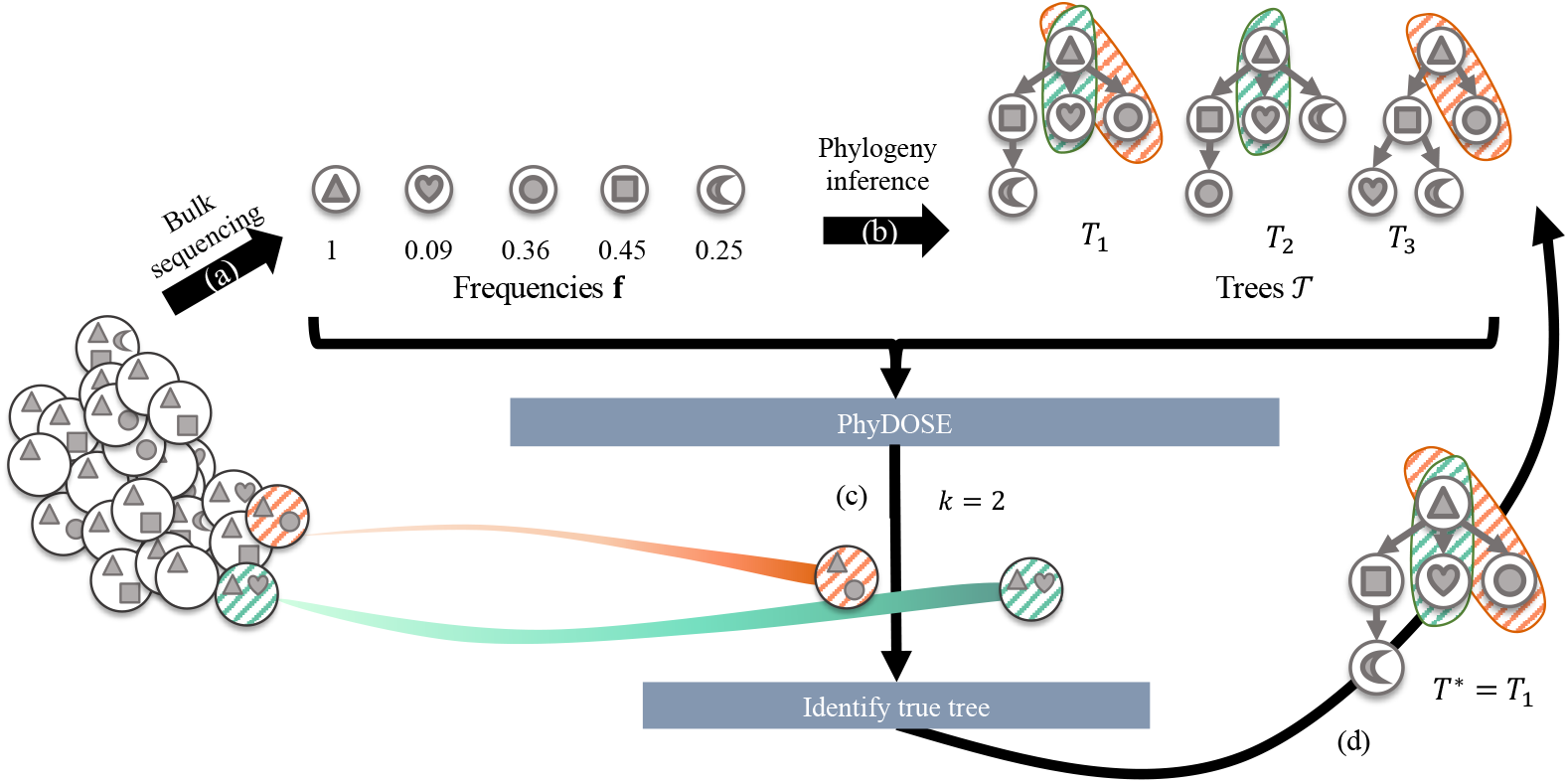
PhyDOSE computes the number of single cells to sequence to identify the true phylogeny. (a) Mutation frequencies **f** obtained from bulk DNA sequencing data. (b) The solution space 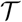 of trees inferred from **f**. We show a distinguishing feature of *T*_1_ (orange and green). (c) For tree *T*_1_, PhyDOSE suggests that *k* = 2 single cells suffice to observe clones that are unique to *T*_1_. (d) In a follow-up SCS experiment we observe *k* = 2 cells, one from the orange clone and one from the green clone. As such, we eliminate trees *T*_2_ and *T*_3_, concluding that phylogeny *T*_1_ is the true phylogeny *T*\*.

Several hybrid datasets have been obtained by performing bulk and single-cell DNA sequencing simultaneously [21,22]. However, there is merit in first performing bulk sequencing to guide follow-up SCS experiments. For instance, several studies first identified a subset of single-nucleotide variants from the bulk data to target in subsequent SCS experiments, thereby reducing costs compared to conventional whole-genome SCS approaches [23–25]. A recently introduced method, SCOPIT, computes how many cells are needed to observe all clones of a tumor, given estimates on the smallest prevalence of a clone as well as the number of clones to detect [26]. The authors provide no guidance on how to obtain these two quantities. Here, we build upon this work by directly incorporating knowledge encoded by the trees 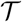 inferred from the initial bulk sequencing data. Indeed, by using data from a SCS experiment we may eliminate trees from 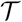 that do not align with the observed clones (Fig 1). In other words, if we observe all clones in a tumor, it is possible to determine the phylogeny of the tumor. However, is it possible to achieve the same goal by observing fewer clones? If so, how many cells are necessary for us to observe the required clones?

We introduce <u>Phy</u>logenetic <u>D</u>esign <u>Of</u> <u>S</u>ingle-cell sequencing <u>E</u>xperiments (PhyDOSE), a method to strategically design a follow-up SCS experiment aimed at inferring the true phylogeny (Fig 1). Given a set 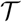 of candidate trees inferred from initial bulk data, we describe how to distinguish a single tree *T* among the rest using features unique to *T*. In particular, if our SCS experiment results in observing cells corresponding to a distinguishing feature of *T*, we may conclude that *T* is in fact the true tree. This means that we can typically identify *T* using only a subset of the clones. To determine the number of cells to sequence, we introduce a probabilistic model that incorporates SCS errors and models successful SCS experiments as a tail probability of a multinomial distribution (Fig 1c). Finally, we reconcile the sampled cells utilizing these distinguishing features to infer the true phylogeny (Fig 1d). We validate PhyDOSE using both simulated data and a retrospective analysis of a leukemia patient that has undergone both bulk and SCS sequencing. We also demonstrate the utility of PhyDOSE by prospectively computing how many cells are needed to resolve the uncertainty in phylogenies of a recent lung cancer cohort. The cost-efficient SCS experiments enabled by PhyDOSE will yield high-fidelity phylogenies, improving downstream analyses aimed at understanding tumorigenesis and developing treatment plans.

## Materials and methods

We introduce <u>Phy</u>logenetic <u>D</u>esign <u>O</u>f <u>S</u>ingle-cell sequencing <u>E</u>xperiments (PhyDOSE), a method to determine the number of single cells to sequence to identify the true phylogeny given initial bulk sequencing data. PhyDOSE is implemented in C++/R and is available at https://github.com/elkebir-group/PhyDOSE. This section describes the various methodological components of PhyDOSE.

### Problem Statement

Let *n* be the number of single-nucleotide variants, or simply *mutations,* identified from initial bulk sequencing data of a matched normal and tumor biopsy sample. For each mutation *i*, we observe the *variant allele frequency* (VAF), i.e. the fraction of aligned reads that harbor the tumor allele at the locus of mutation *i*. Specialized methods exist that combine copy number information and VAFs to infer a *cancer cell fraction* f¿ for each mutation *i*, which is the proportion of cells in the tumor biopsy that contain at least one copy of the mutation [3,27–29]. Here, we refer to cancer cell fractions as *frequencies*. Typically, phylogenies 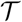 inferred by current methods from frequencies **f** = [*f_i_*] adhere to the infinite sites assumption. That is, each mutation i is introduced exactly once at vertex *v_i_* and never subsequently lost.

When we sequence a single cell from the same tumor biopsy, assuming no errors, we identify a clone of the tumor. In other words, we observe a set of mutations that must form a connected path in the unknown true phylogeny *T*\*. By repeatedly sequencing single cells until we observe all clones in the tumor, we will have observed all root-to-vertex paths of *T*\*, thus identifying tree *T*\* itself. We assume that (i) the true unknown phylogeny *T*^*^ is among the trees in 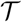 and that (ii) mutations among single cells that we sample from the tumor biopsy follow the same distribution as **f**. These assumptions are important for the mathematical derivation of PhyDOSE but it is typical for violations to occur in practice. Through simulations, we explore the impact of violating these assumptions and show that our approach is robust to many realistic scenarios.

This leads to the following question and problem statement with respect to these two assumptions. How many single cells do we need to identify *T*\* with confidence level *γ*?

#### Problem 1

(SCS Power Calculation (**SCS-PC**)). Given a set 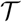 of candidate phylogenies, frequencies **f** and confidence level *γ*, find the minimum number *k*\* of single cells needed to determine the true phylogeny *T*\* among 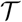 with probability at least *γ*.

Clearly, we do not know which phylogeny in 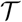 is the true underlying phylogeny *T*\* of the tumor. Thus, we consider a slightly different problem: In the *T*-SCS-PC problem (defined formally at the end of the section), we are given an arbitrary phylogeny 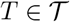 and want to perform a similar power calculation when conditioning on *T* being the true phylogeny. By solving the T-SCS-PC problem for all trees 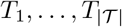, we obtain the numbers 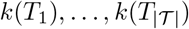 of single cells needed for each tree. As *T*\* is in 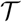, the maximum number among 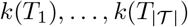 is an upper bound on the number of required SCS experiments to identify *T*\* with probability at least *γ*. To solve the T-SCS-PC problem, we need to reason for which SCS experiments we can conclude that *T* is the true phylogeny.

Observe that each tree *T* in 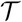 describes a unique set of clones, corresponding to the sets of mutations encountered in all root-to-vertex paths of *T* (Fig 1). Thus, if we observe all clones of a phylogeny *T* in our SCS experiments, we may conclude that T is the true phylogeny. What is the probability of doing so? To answer this question, we must compute the prevalence of each clone in the tumor biopsy.

For phylogenies that adhere to the infinite sites assumption, the *prevalence* **u**(*T*, **f**) = [*u_i_*] of the clones in the tumor biopsy are uniquely determined by the phylogeny *T* and frequencies **f** as

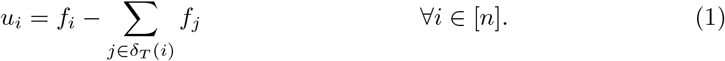

where *δ_T_*(*i*) is the set of children of the node where mutation i was introduced [9]. Tumor phylogeny inference methods guarantee that the inferred phylogenies 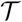 from frequencies **f** have clonal prevalence **u**(*T*, **f**) = [*u_i_*] that are nonnegative and that
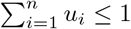, where the remainder 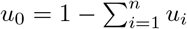 is the prevalence of the normal clone. Thus, conditioning on a phylogeny *T* and frequencies **f**, sequencing one cell from the tumor will lead us to observe one of the *n* +1 clones of *T* with probabilities (*u*_0_,…, *u_n_*). In other words, the outcome of this SCS experiment with one cell is a draw from the categorical distribution Cat(*u*_0_,…,*u_n_*). The possible outcomes of a SCS experiment composed of *k* cells thus follow a multinomial distribution Mult(*u*_0_,…, *u_n_*). Thus, the probability of observing all tumor clones of *T* in such a SCS experiment with *k* cells corresponds to the tail probability of the multinomial where each of the *n* tumor clones is observed at least once.

The corresponding *power calculation* is to determine the smallest number for *k* where the tail probability is greater or equal to the confidence level *γ*. Note that this power calculation for observing all clones has been previously introduced [26].

Importantly, in many cases we need not observe all clones of *T* to distinguish *T* from the remaining phylogenies 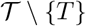 (Fig 2). This means that we may conclude that *T* is the true phylogeny with a SCS experiment with fewer cells. To formalize this notion, we start by defining a featurette.

**Fig 2.**
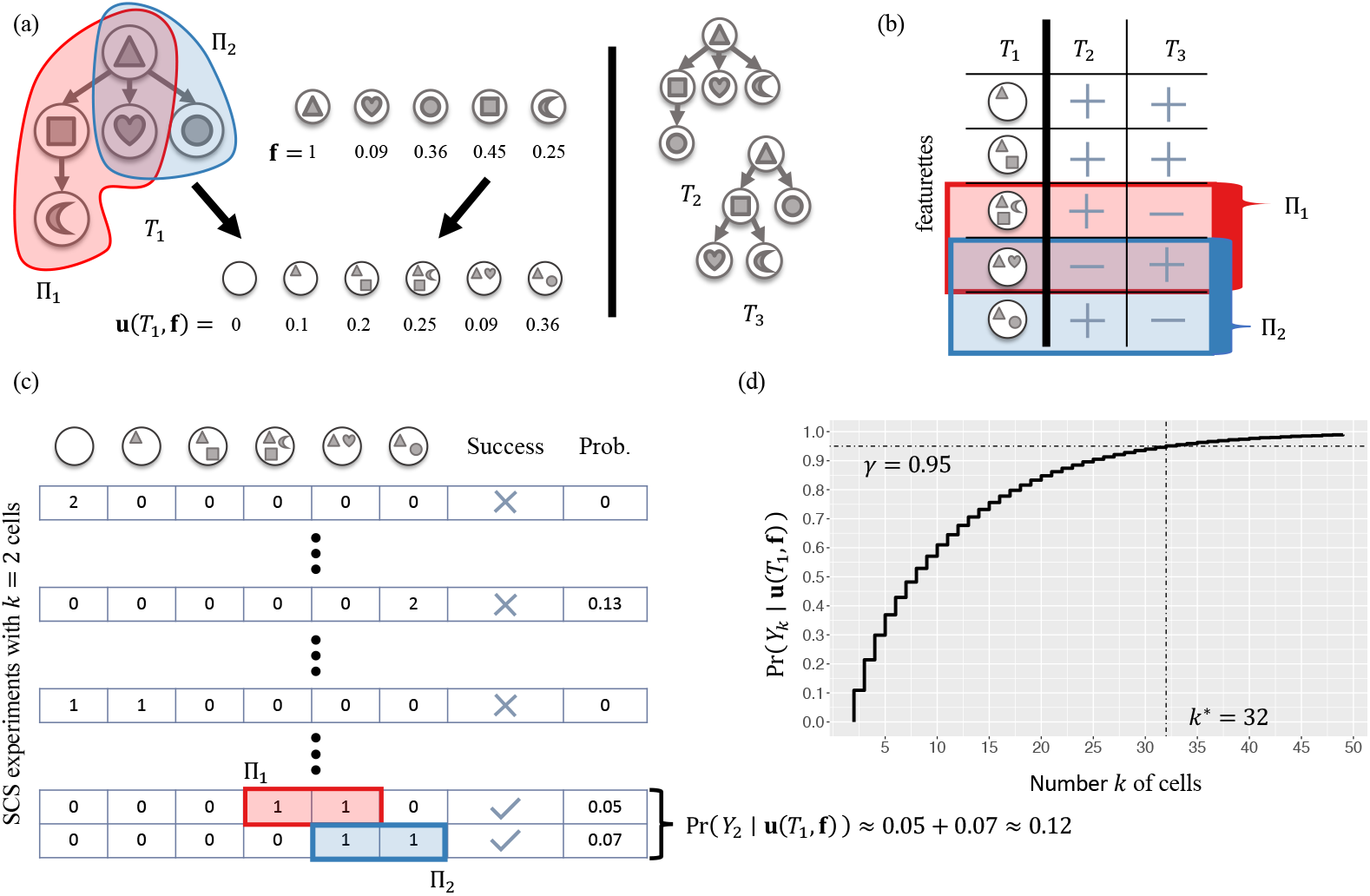
The SCS Power Calculation for Phylogeny *T*(*T*-SCS-PC) problem. (a) We are given frequencies **f** and a tree *T*_1_ that we want to distinguish from the other trees {*T*_2_,*T*_3_}. The pair (*T*_1_, **f**) uniquely determine clonal prevalence **u**(*T*_1_, **f**). (b) Featurettes of *T*_1_ correspond to root-to-vertex paths, yielding distinguishing features Π_1_ and Π_2_, each with one featurette absent in *T*_2_ and another absent in *T*_3_. (c) With *k* = 2 cells, we must observe clones from either Π_1_ or Π_2_ for a successful outcome, resulting in probability Pr(*Y*_2_ | **u**(*T*_1_, **f**)) ≈ 0.12. (d) To increase this probability to *γ* = 0.95, we need *k*\* = 32 cells.

#### Definition 1. A *featurette* τ is a subset of mutations

We say that a featurette *τ* is *present* in a phylogeny *T* if the nodes/mutations of *τ* form a connected path of *T* starting at the root node, otherwise we say that *τ* is *absent* in *T*. The same featurette, however, may be present in more that one phylogeny. Thus, multiple featurettes may be required to distinguish a phylogeny *T* from the remaining phylogenies 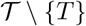.

#### Definition 2.

A set Π of featurettes is a *distinguishing feature of T* if (i) for all featurettes *τ* ∈ Π it holds that *τ* is present in *T*, and (ii) for each remaining phylogeny 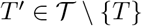 there exists a featurette *τ*’ ∈ Π where *τ*’ is absent in *T*’.

Thus, a SCS experiment where we observe one cell from each clone of a distinguishing feature Π of *T* enables us to conclude that phylogeny *T* is the true phylogeny. As discussed, every phylogeny *T* has a *trivial distinguishing feature*, which is composed of all featurettes present in *T*. Moreover, *T* may have multiple distinguishing features. Therefore, we must consider the complete set of all distinguishing features, which we call the distinguishing feature family.

#### Definition 3.

The set 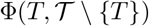 composed of all distinguishing features of *T* with respect to 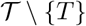 is a *distinguishing feature family of T*.

Let (*c*_0_,…, *c_n_*) be the outcome of a SCS experiment of *k* cells, where *c_i_* ≥ 0 is the number of cells observed of clone *i* and 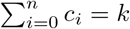. This experiment is *successful* if, among the *k* sequenced cells, we observe the clones of at least one distinguishing feature 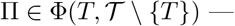 i.e. *c_i_* > 0 for all clones i in some distinguishing feature 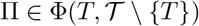. As discussed, conditioning on frequencies **f** and *T* being the true phylogeny, outcomes (*c*_0_,…, *c_n_*) of SCS experiments of *k* cells follow a multinomial distribution Mult(*k, u*_0_,…, *u_n_*) where **u**(*T*, **f**) = [*u_i_*] is defined as in (1). Let *Y_k_* denote the event of a successful outcome. We are interested in computing the probability Pr(*Y_k_* | **u**(*T*, **f**)), which equals the sum of the probabilities of all successful outcomes. More specifically, we want to determine the smallest number *k*^*^ of single cells to sequence such that Pr(*Y*_*k*\*_ | **u**(*T*, **f**)) is at least the prescribed confidence level *γ* (Fig 2).

#### Problem 2

(SCS Power Calculation for Phylogeny *T* (*T*-SCS-PC)). Given a set 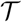 of candidate phylogenies and a phylogeny 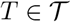, frequencies **f** and confidence level *γ*, find the minimum number *k*\* of single cells needed such that Pr(*Y_k_* | **u**(*T*, **f**)) ≥ *γ*.

.In Section A.1 in S1 Text, we prove that the above problem is NP-hard.

#### Theorem 1.

*T*-SCS-PC is NP-hard.

### Multinomial Power Calculation

To solve the *T*-SCS-PC problem, it suffices to have an algorithm that computes Pr(*Y_k_* | **u**(*T*, **f**)), which is the probability of concluding that *T* is the true phylogeny. Using this algorithm we identify *k*\* by starting from *k* = 0 and simply incrementing *k* until the corresponding probability Pr(*Y_k_* | **u**(*T*, **f**)) exceeds the prescribed confidence level *γ*. In the following, we describe how to efficiently compute Pr(*Y_k_* | **u**(*T*, **f**)).

Recall that the outcome of a SCS experiment composed of *k* cells corresponds to a vector **c** = [*c_i_*], where *c_i_* ≥ 0 is the number of cells that we observe from clone *i* and 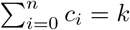. In a successful outcome **c** we observe at least one cell for each featurette in at least one distinguishing feature 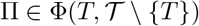, where 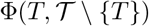 is the distinguishing feature family. For brevity, we will write Φ rather than 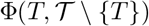.

Let **c**(Π, *k*) denote the set of all outcomes where we observe at least one cell for each featurette in a distinguishing feature Π — i.e. 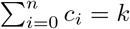, and for all *i* ∈ {0,…, *n*} it holds that *c_i_* > 0 if clone *i* is a featurette in Π and *c_i_* ≥ 0 otherwise. The set **c**(Φ, *k*) of successful outcomes is defined as the union 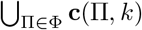. The probability of any SCS outcome **c** = (*c*_0_,…, *c_n_*) is distributed according to Mult(*k*, **u**(*T*, **f**)). Since successful outcomes enable us to conclude that T is the true phylogeny, we have

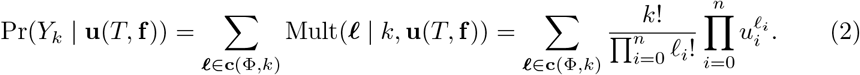

If there is only one distinguishing feature Π, i.e. Φ = {Π}, then the desired probability is a standard tail probability of the multinomial where we sum up the probabilities of outcomes **c**(Π, *k*) = [*c_i_*] such that 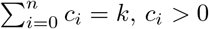 if clone *i* is a featurette of Π and *c_i_* ≥ 0 otherwise. A fast calculation of this tail probability was developed using a connection to the conditional probability of independent Poisson random variables [26,30]. If there are multiple distinguishing features but they are pairwise disjoint — i.e. no two distinct distinguishing features share the same featurette — then we simply have

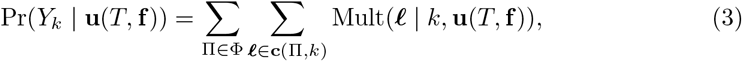

and we can apply the fast computation [26] to obtain each independent tail probability. However, the equality in the above equation does not hold if the family Φ is composed of distinguishing features with overlapping featurettes. Incorrectly applying this equation will lead us to overestimate the value of *k*\*. Since single-cell sequencing is expensive, overestimating the number of cells to sequence in a SCS experiment can be costly and unnecessary. One naive way would be to simply brute force all (*n* + 1)^*k*^ SCS outcomes, but this will not scale. Instead, to calculate Pr(*Y_k_* | **u**(*T*, **f**)) exactly, we propose to use the inclusion-exclusion principle as follows.

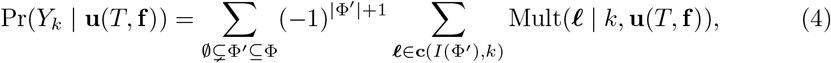

where *I*(Φ’) is the set of all featurettes in Φ’, i.e. 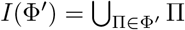 (Fig 3a).

**Fig 3.**
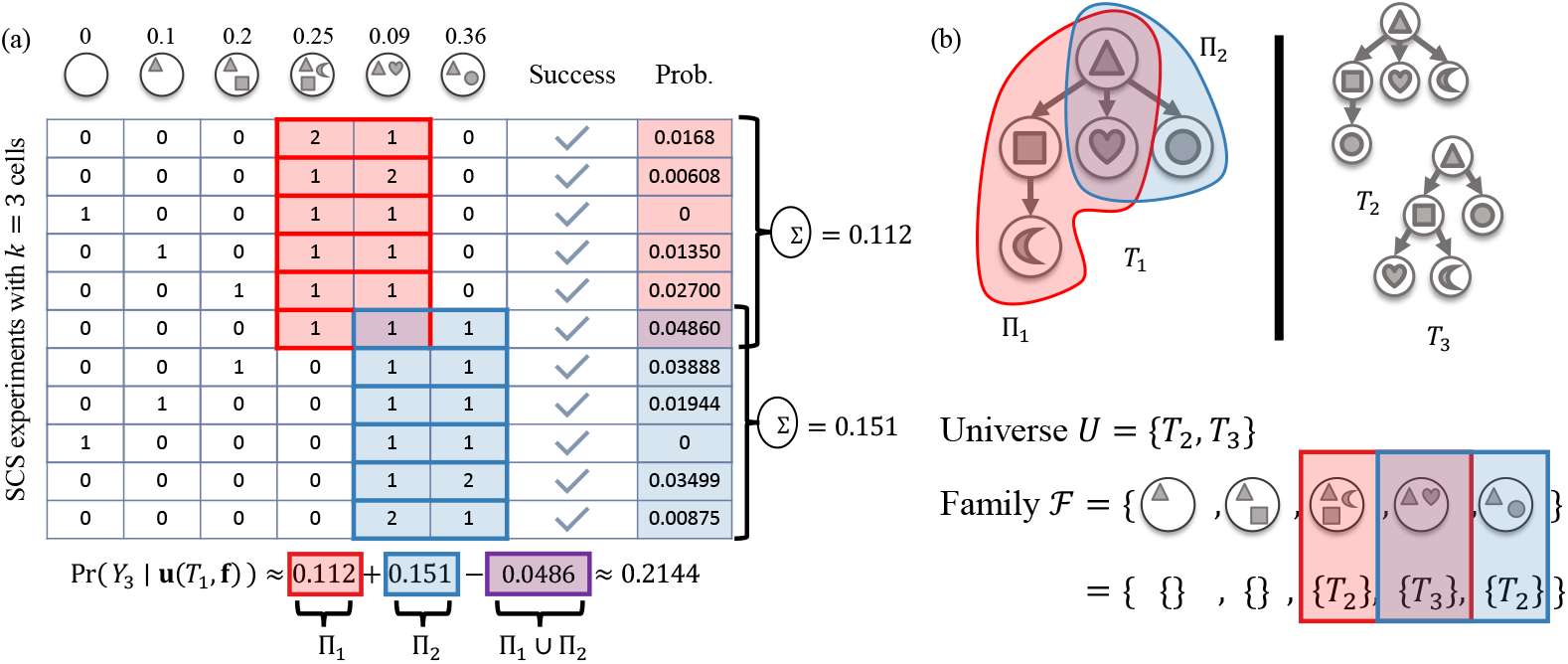
PhyDOSE implementation details. (a) To account for minimal distinguishing features that share featurettes, we use the inclusion-exclusion principle to compute Pr(*Y_k_* | **u**(*T*, **f**)). Here, Π_1_ (red) and Π_2_ (blue) share a featurette (with ‘triangle’ and ‘heart’ mutations). (b) To enumerate the set Φ* of minimal distinguishing features of *T*_1_, we reduce the problem to Set Cover and repeatedly identify minimum covers. Here, the universe *U* is composed of trees {*T*_2_, *T*_3_} and there is a subset in 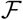 for each featurette *τ* of *T*_1_ composed of the trees where *τ* is absent.

Thus, we need to compute 2^|Φ|^ — 1 tail probabilities, which each can be done using the fast calculation in SCOPIT [26].

In the worst case, Φ has *O*(2^*n*^) distinguishing features resulting in *O*(2^*n*^) tail probabilities. We now describe one final optimization that will significantly reduce the number of required computations. This is based on the following observation.

#### Observation 1.

If Π is a distinguishing feature of *T* then for all featurettes *τ* present in *T* it holds that Π ∪ {*τ*} is a distinguishing feature of *T*.

This means that distinguishing features in Φ form a partially ordered set under the set inclusion relation. We call a distinguishing feature Π *minimal* if there does not exist another distinguishing feature Π’ ∈ Φ that is a proper subset of Π, i.e. 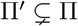.

A direct consequence of Observation 1 is that the outcome of an SCS experiment is successful when we observe all featurettes of a distinguishing feature Π, and remains so even if we observe additional featurettes *τ*’ ∉ Π.

As such, successful outcomes w.r.t. Φ equal those w.r.t. the set Φ* of all minimal distinguishing features of *T*.

#### Observation 2.

It holds that **c**(Φ*, *k*) = **c**(Φ, *k*).

Therefore, it suffices to restrict our attention to only Φ^*^ rather than the complete family Φ when computing Pr(*Y_k_* | **u**(*T*, **f**)) using (4). Section B.1 in S1 Text describes how to find Φ* by reducing the problem to that of finding all minimal set covers, which we solve in an iterative fashion using integer linear programming.

### Consideration of SCS Error Rates

One current challenge with SCS is that the false negative rate per site is quite high with typical rates up to 0.4 for the commonly used multiple displacement amplification (MDA) method [31]. On the other hand, current false positive rates are low and are typically less than 0.0005 for MDA-based whole-genome amplification [31]. A *false negative* is defined as not observing a mutation that is present in the cell. A *false positive* occurs when we observe the presence of a mutation that did not occur in that cell.

With PhyDOSE, we propose one possible method for incorporating the false negative rate *β* when it is known. Specifically, sampled cells follow a categorical distribution **u** = [*u*_0_,…, *u_n_*] when conditioned on tree *T*. Hence, the probability of sampling a cell from clone *i* equals *u_i_*. True positives, i.e. correctly observing a mutation in a clone, follow a Bernoulli distribution with parameter 1 – *β*. To observe a featurette/clone *i* that has *n_i_* mutations and a prevalence of *u_i_*, we thus need to have *n_i_* true positives. In other words, assuming independence among mutations, we require *n_i_* successful draws from a Bernoulli distribution parameterized by 1 — β. As such, we derive new clonal prevalence 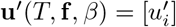 from **u**(*T*, **f**) = [*u_i_*]. Additionally assuming independence between the events of a cell being sampled from clone *i* and the absence of false negatives, we set 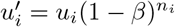 where *n_i_* is the number of mutations in featurette/clone *i*. We set 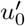 to be equal to 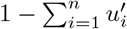. This adjustment results in a reduction of the clonal prevalence and ultimately increases the value of *k*\*. The issue of false positives is less serious as error rates are low enough to be negligible.

### Prioritizing Candidate Trees Post SCS Experiment

The final step is to prioritize candidate trees after performing a SCS experiment with the number *k*\* of cells computed by PhyDOSE. To this end, we compute the *support* of each tree 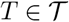. Intuitively, support(*T*) is the number of cells that support the conclusion that *T* is the actual phylogeny. Formally, we say that a distinguishing feature Π of a tree *T* is *observed* if each featurette of Π is observed in at least one cell.

Using this, we define support(*T*) as the number of cells that correspond to featurettes of an observed distinguishing feature Π of *T*. Per Observation 1, it suffices to restrict our attention to the set Π^*^ of minimal distinguishing features.

There are two outcomes of a SCS experiment with *k*\* cells. Either there is no tree 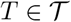 with non-zero support or there are one or more trees with non-zero support. In the former case, the SCS experiment has failed, which is expected to occur with probability 1 – *γ*. In the latter case, which may occur in the presence of false negatives and false positives, we return the set of trees with maximum support.

Alternatively, we may use existing methods that infer tumor phylogeny from SCS data [14,16,32] or a combination of SCS and bulk data [20,33].

## Results

In this section, we demonstrate the application of PhyDOSE to simulated and real data. We begin by validating our method using simulated data. Next, we provide retrospective results for a leukemia patient [23] and an acute myeloid leukemia cohort [34] where both bulk and single-cell DNA sequencing have been performed [23]. Finally, we use PhyDOSE to perform a prospective analysis to determine the required number of single cells to identify the true phylogeny in a non-small cell lung cancer patient cohort [3].

### Simulations

#### Design

To assess the performance of PhyDOSE, we generated simulated data where the ground truth tree *T*\* is known. Given a fixed number *c* of clones and *n* mutations, we first generated a ground truth tree *T*\* with *c* vertices uniformly at random using Prufer sequences [35] and randomly distributed the *n* mutations to the *c* clones while ensuring that every clone had at least one mutation. Next, we generated clonal prevalences **u** = [*u_i_*] by drawing from a symmetric *n* + 1-dimensional Dirichlet distribution with concentration parameter 0.2. We used rejection sampling to ensure that each clonal prevalence *u_i_* was at least 0.05. Let *σ*(*i*) be the set of clones that contain mutation *i*. We generated frequencies **f** = [*f_i_*] by setting 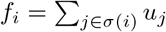 for each mutation *i* ∈ { 1, …, *n*}.

To validate our method using *in silico* SCS experiments, we introduced errors related to single-cell sequencing by varying false negative rates *β* ∈ {0, 0.2} and doublet rates *δ* ∈ {0, 0.1}. We generated, for each simulation instance, 10, 000 single cells sampled either under the specified false negative rates *β* and doublet rates *δ* according to the bulk clonal prevalence **u**. Additionally, we varied single-cell clonal prevalences from the bulk clonal prevalences by resampling **û** ~ D_IR_(λ**u**). We tuned the parameter λ so that the clonal prevalence varied by an absolute average of 5% from the clones of the ground truth tree *T*\*, which resulted in λ = 2000. We performed further sensitivity analysis (S2 Fig) on the robustness to clonal prevalence distortion by investigating the impacts of setting λ = 50, equating to an absolute average difference of 20% between the bulk and single cell clonal prevalence.

In total, we generated 100 simulation instances under eight varying conditions as specified in Table 1. We used the SPRUCE algorithm to enumerate the set 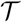 of trees given frequencies **f** [9]. In order to consider violations of the PhyDOSE assumption that the true tree is among the candidate set, for simulation conditions ‘b’ (sim1b, sim2b and sim3b), we randomly sampled 10% of the trees outputted by SPRUCE [9]. For sim2c, we introduce a clonal prevalence distortion of 20%.

**Table 1.** Simulation conditions. We generate simulated data under eight conditions with 100 instances each. These conditions have varying subsets of candidate trees, number of clones, number of mutations per clone, clonal prevalence distortions, false negative and doublet rates.

| ID | % of Trees | Clones | Mutations | Prevalence Noise | FNR $\beta$ | Doublet $\delta$ |
| --- | --- | --- | --- | --- | --- | --- |
| sim1a | 100% | 7 | 7 | 0% | 0 | 0 |
| sim1b | 10% | 7 | 7 | 0% | 0 | 0 |
| sim2a | 100% | 7 | 7 | 5% | 0 | 0 |
| sim2b | 10% | 7 | 7 | 5% | 0 | 0 |
| sim2c | 100% | 7 | 7 | 20% | 0 | 0 |
| sim3a | 100% | 7 | 7 | 5% | 0.2 | 0.1 |
| sim3b | 10% | 7 | 7 | 5% | 0.2 | 0.1 |
| sim4a | 100% | 10 | 100 | 5% | 0.2 | 0.1 |

For sim4a, we first used PyClone [36] to cluster the 100 mutations before enumerating the set.

#### Results

We now compare the number *k*\* of single cells computed by PhyDOSE to the naive method of requiring all clones to be observed. In addition, we perform *in silico* SCS experiments using PhyDOSE’s computed number *k*\* to assess its accuracy in light of varying sequencing and sampling errors.

The number 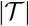 of trees in our simulations ranged from 3 to 506 for the full enumerated set of trees (simulation conditions ‘a’) with a median of 59 trees (Fig 4a). For the set of downsampled trees (simulation conditions ‘b’), the number of trees ranged from 2 to 50 with a median of 7 trees (Fig 4a).

**Fig 4.**
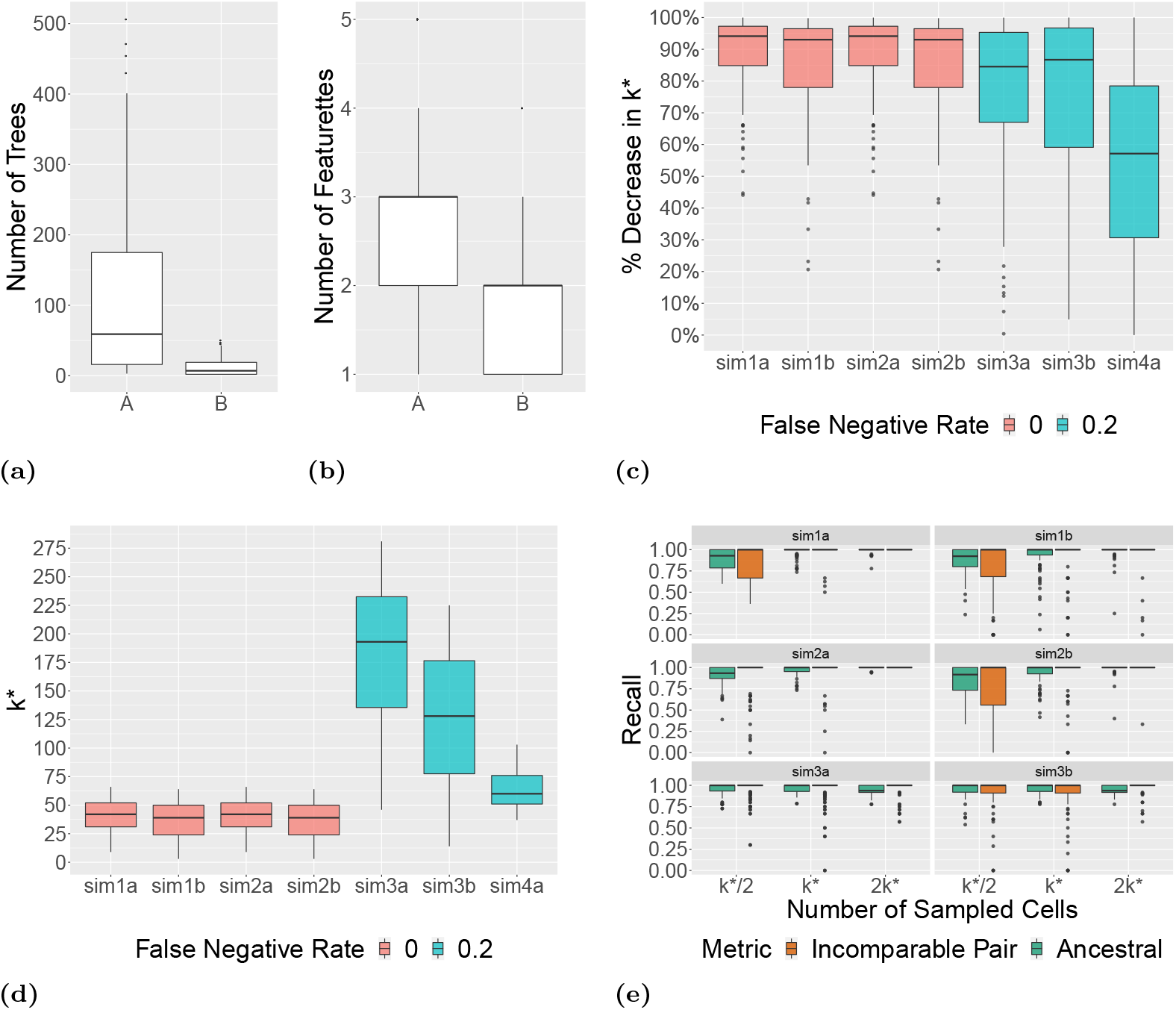
Simulations demonstrate that PhyDOSE’s calculated number of single cells resolves tree ambiguity in bulk sequencing data. We used confidence level *γ* = 0.95 to determine the number *k*\* of single cells to sequence. (a) Number 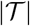 of trees output by SPRUCE [9]. (b) Number |Π| of featurettes among minimal distinguishing features Φ*. (c) Percent decrease in *k*\* when utilizing PhyDOSE instead of the naive method. (d) Number *k*\* of cells identified by PhyDOSE. (e) Recall metrics of the tree inferred by SPhyR [16] by randomly sampling *k*\*/2, *k*\* and 2*k*\* simulated single cells.

We ran PhyDOSE to identify the minimal distinguishing feature family Φ* for each tree in each simulation instance. This yielded a single minimum distinguishing feature in each case for the simulations instances with fully enumerated tree sets, whereas for the downsampled tree set the number of distinguishing features was between 1 and 6 with a median of 2. Fig 4b shows the number of featurettes in each minimal distinguishing feature identified by PhyDOSE, ranging from 1 to 5 with a median of 3 for conditions ‘a’ and 2 for conditions ‘b’. Importantly, this number is smaller than the total number of 5 featurettes. As such, running PhyDOSE resulted in a median reduction of ~ 88% over all simulations in the follow-up experimental design compared to the naive approach of requiring all featurettes/clones to be observed (Fig 4c,d).

In particular, with a confidence level of 7 = 0.95, PhyDOSE computed a median number of *k*\* = 41 cells compared to *k*\* = 544 cells computed by the naive method (Fig 4d). With a false negative error rate *β* = 0.2, PhyDOSE computed *k*\* = 137 cells compared to *k*\* = 720 cells by the naive method (Fig 4d). Thus, with increasing false negative rates *β* ∈ {0, 0.2} we observed that (i) PhyDOSE continued to outperform the naive method and (ii) more cells were needed to identify *T*\*.

Next, we assess the accuracy of PhyDOSE’s *k*\* value for varying single-cell sequencing error conditions. To do so, we ran our approach for prioritizing candidate trees as well as SPhyR [16] on sampled single cells. For the former, we performed 100 experiments for each simulation instance, reporting the number of experiments that successfully recovered the ground truth tree *T*\*. To test the precision of *k*\* we also considered the performance of SPhyR when randomly sampling half and double the *k*\* cells determined by PhyDOSE. S3 Fig shows that the prioritization approach worked well in the case the candidate tree set contains the ground truth tree (median of 89% success rate for conditions sim1a, sim2a and sim3a), while performance dropped when downsampling candidate trees (median success rate of 0% for sim1b, sim2b and sim3b).

Nevertheless, for both sets of simulation conditions, SPhyR [16] was able to identify the true tree in the majority of cases after sampling PhyDOSE computed number *k*\* of cells. In the cases where the estimated tree *T* and true tree *T*\* were different, SPhyR still performed well in accordance with two commonly-used tree distance metrics, ancestral and incomparable pair recall. *Ancestral pair recall* is defined as 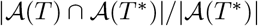 where 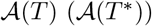 is the set of ordered pairs of mutations that occur on distinct edges of the same branch of *T* (*T*\*). *Incomparable pair recall* is defined as 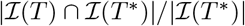 where 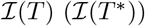 is the set of unordered pairs of mutation that occur on edges in distinct branches in *T* (*T*\*). The median of both metrics is 1 when sampling *k*\* cells (Fig 4e). Additionally, we found greater gains in performance between sampling *k*\* cells versus *k*\*/2 cells than sampling 2*k*\* cells versus *k*\* cells, with this trend less pronounced in sim3a and sim3b.

Next, we assessed the impact of clonal prevalence distortion between bulk and single cell data. We found PhyDOSE to be robust to random clonal prevalence noise between bulk and single-cell sequencing as evidenced by a drop of only 1% in the median percentage of successful *in silico* SCS experiment when there is no downsampling of trees (S2 Fig). Additionally, the recall performance metrics (Fig 4e, S2 Fig) are also not substantially different between sim1a versus sim2a and sim2c. We attribute this to the fact that PhyDOSE’s use of distinguishing features critically relies on the clonal prevalence of a few key clones. Furthermore, only when *u_i_* > *û_i_* for such a key clone i will PhyDOSE will underestimate *k*(*T*), yielding conservative estimates otherwise.

Moreover, we note a drop in the median ancestral pair recall between *k*\* and 2*k*\* when introducing false negatives and doublets in sim3a and sim3b (Fig. 4e). We attribute this to utilizing the 100th percentile for *k*\*, setting a confidence level of 0.95 and adjusting for *β* = 0.2 leading to highly conservative estimates on the number of cells to sequence. Therefore we do not anticipate a significant change in performance to be evident within these three particular sampling scenarios (*k*\*/2, *k*\* and 2*k*\*). We note that there is similar performance between sim2a/sim2b and sim3a/sim3b at *k*\*, which likely demonstrates the benefit of adjusting PhyDOSE for the false negative rate. Although PhyDOSE nor SPhyR [16] explicitly account for doublets, tree recall metrics remain high.

Even when the candidate set is significantly downsampled prior to running PhyDOSE, SPhyR still returned trees with high recall performance. There was a small increase in variation between simulations with the full set of candidate trees and the reduced set (Fig 4e). For sim1 and sim2, the distribution of *k*\* overlaps substantially with a median per replication difference in *k*\* of 1 (IQR: 0-3). Therefore, similar performance is expected in sim1a vs sim1b and sim2a vs sim2b. The median per replication difference in *k*\* between sim3a and sim3b was 55 (IQR: 35.5-82). Thus, elevated variation in the recall metrics is expected in those replications where *k*\* is significantly underestimated.

S4 Fig shows PhyDOSE performance with mutation clusters inferred by PyClone [36]. We additionally included the clustered pair recall in our analysis where clustered pair recall is 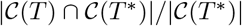, where 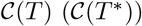 is defined as the set of unordered pairs of mutations that are introduced on the same edge in *T* (*T*\*). At *k*\* cells, the median ancestral pair recall was 0.96, the incomparable pair recall was 0.86 and the clustered pair recall was 0.94, showing a reduction in performance from the first three simulations due to the additional errors introduce by PyClone [36].

In summary, our simulations demonstrate that PhyDOSE’s distinguishing feature analysis results in significantly fewer cells to sequence than the naive approach without a subsequent loss in power to identify the true phylogeny. Moreover, we find that PhyDOSE is robust to increasing values of false negatives, doublets and clonal prevalence noise that are typical to real data as well to the case when 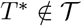.

#### Running time

Performing the calculation of *k*\* is fast [26] when |Φ| = 1 and the median of |Φ| in our simulations was 1. Therefore, the main bottleneck in PhyDOSE is the determination of Φ for each tree in the candidate set as this involves iteratively solving an ILP for each tree until all minimal distinguishing features are found. Note that this step is embarrassingly parallelized because Φ is computed independently for each 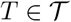. To explore a practical upper bound on 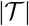, we calculated the runtime in seconds for each simulation instance when Φ is solved sequentially for each 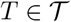. To additionally explore how PhyDOSE scales with 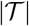, we generated an additional simulation set of with 10 mutations and no mutations clusters and set a two-hour time limit. The largest sized set was 40,089 trees with a median of 112 trees. The results are displayed in S5 Fig. Simulations where 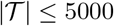 completed within the two hour time limit when solved sequentially, suggesting an upper bound on 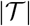 to use in practice without parallelization. The mean runtime per tree grows linearly with 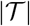, thus providing a means of estimating total runtime with parallelization and/or large sized inputs.

### Retrospective Analysis of an Acute Lymphoblastic Leukemia Patient

We considered a cohort of six childhood acute lymphoblastic leukemia (ALL) patients whose blood was sequenced using bulk and targeted single-cell DNA sequencing [23]. The number of sequenced single cells per patient varied between 96 and 150. To validate our approach, we used PhyDOSE to calculate the number *k*(*T*\*) of cells needed to identify the true phylogeny *T*\* that is consistent with both data types, thereby retrospectively determining whether fewer single cells suffice to determine *T*\*, decreasing the cost of replicate experiments. In addition, we assessed whether the calculated number *k*(*T*\*) yielded *T*\* using *in silico* SCS experiments.

Due to the absence of published copy-number aberration information for this dataset, we focused our attention on patient 2 whose single-cell phylogeny adhered to the infinite sites assumption and the variant allele frequencies suggested the absence of copy-number aberrations (as detailed in Section C in S1 Text). For this patient, 16 autosomal mutations in 115 cells were sequenced [23]. We note that the authors had no knowledge of the number of cells that would suffice to infer the tumor phylogeny of the patient. Using the infinite sites assumption and assuming the absence of copy-number aberrations, we define the cancer cell fraction, or frequency *f_i_* of each mutation *i* in the bulk data as 2 · VAF(*i*). We define the *SCS mutation frequency* as the fraction of single cells that harbor the mutation. Strikingly, there is a clear correlation between the bulk and SCS mutation frequencies, supporting PhyDOSE’s first assumption (Fig 5a). We excluded mutation *CMTM8* because of a notable discrepancy in frequencies (0.4 in bulk vs. 0.2 in SCS). Using SPRUCE [9], we enumerated the set 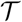 of trees from the bulk data, yielding over 2.5 million trees. This number is mainly driven by 3 mutations *(ATRNL1, LINC00052* and *TRRAP*) with a VAF less than 0.05. Excluding these 3 mutations resulted in a more tractable number of 2576 trees. We note that in practice we may similarly exclude mutations because of very low VAFs or less importance in downstream analyses. Fig 5b shows the single tree 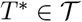 that was consistent with the cleaned single-cell data, supporting PhyDOSE’s second assumption.

**Fig 5.**
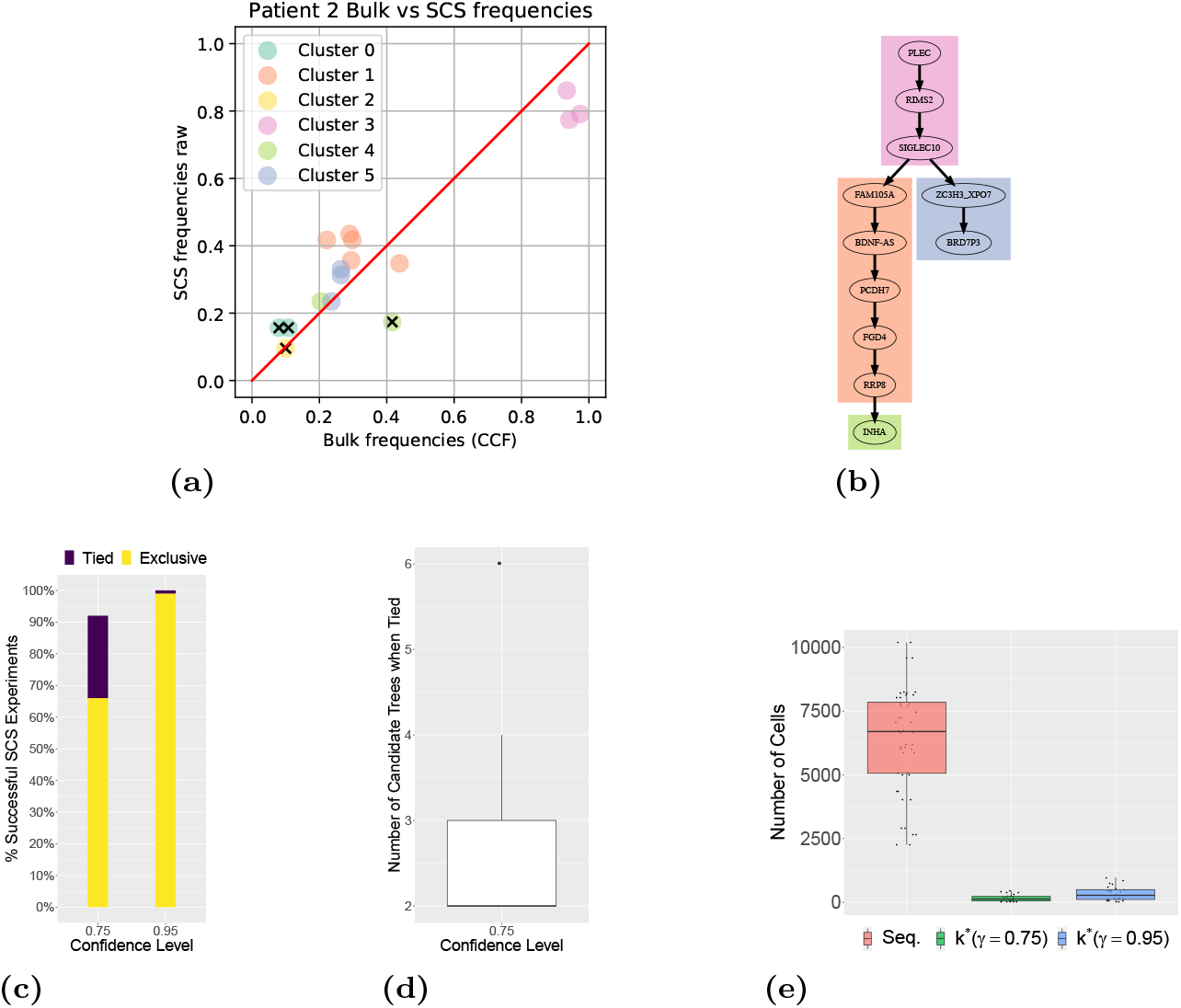
Retrospective analysis of ALL patient 2 [23] and AML cohort [34] demonstrates that fewer cells suffice for replication. Panels (a)-(d) consider ALL patient 2 [23] and panel (e) considers the AML cohort [34]. (a) There is a strong correlation between bulk and single-cell mutation frequencies. Colors indicate mutation clusters from SCS data and excluded mutations are indicated by ‘x’. (b) Phylogeny T* that is consistent with the SCS and bulk data. (c) Percent of successful outcomes in 100 *in silico* SCS experiments, obtained by sampling from the 115 sequenced cells without replacement following PhyDOSE’s calculated number *k*(*T*\*) of cells (103 for *γ* = 0.95 and 50 for *γ* = 0.75). Exclusive outcomes (yellow) uniquely identified *T*\* whereas tied outcomes (purple) yielded a small set of candidate phylogenies that include *T*\*. (d) Number of candidate phylogenies in the case of ties. (e) The distribution of PhyDOSE’s *k*\* for *γ* ∈ {0.75, 0.95} of all patients in the AML cohort with 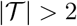 as well as the number of cells that were originally sequenced.

We ran PhyDOSE using varying confidence levels *γ* ∈ {0.75,0.95} and an estimated false negative rate of *β* = 0.2 reported by the authors [23]. PhyDOSE calculated that *k*(*T*\*) = 103 cells suffice to identify T* with confidence level *γ* = 0.95. Indeed, performing 100 *in silico* SCS experiments, by sampling *k*(*T*\*) cells among the 115 sequenced cells without replacement, yielded a success rate of 99% (Fig 5c).

To reduce costs, we explored what would have happened retrospectively with a lower confidence level *γ* of 0.75. PhyDOSE calculated that *k*(*T*\*) = 50 cells are needed for *γ* = 0.75, which is a significant cost savings over *γ* = 0.95. Performing 100 *in silico* SCS experiments yielded a success rate of uniquely identifying *T*\* of 66%, which was lower than the expected rate of 75%. Furthermore, we noted that in an additional 26% of experiments the correct phylogeny *T*\* was among the trees with the highest overall support (Fig 5c). The number of trees in the tied set of successes varied from 2 to 6 (S7d Fig), showing that although PhyDOSE did not uniquely identify the tree, it was able to significantly reduce the original set of 2576 trees (S7 Fig and S8 Fig).

In summary, this retrospective analysis shows that the true tree for patient 2 could have been identified confidently with fewer cells than the 115 cells initially sequenced [23]. With a lower confidence level *γ*, PhyDOSE computes that far fewer cells are required, significantly reducing costs but at the expense of a lower success rate of uniquely identifying the true phylogeny. Nevertheless, the resulting SCS experiment will eliminate a large fraction of the original set of candidate phylogenies due to the incorporation of distinguishing features in the PhyDOSE power calculation.

### Retrospective Analysis of an Acute Myeloid Leukemia Cohort

In [34], high-throughput targeted microfluidic single-sequencing was performed on a cohort of 77 patients with acute myeloid leukemia (AML). The authors additionally performed bulk sequencing in order to confirm the presence of a mutation in the single-cell data. We note that the authors restricted their analysis to somatic mutations (SNVs and indels) that did not occur in regions affected by additional copy number aberrations.

Here, we utilized the published bulk sequencing VAFs of the SNVs in each patient, eliminating any mutations not detected via bulk sequencing, to enumerate a set of candidate trees using SPRUCE [9]. The median number of mutations per patient was 4 (IQR: 3-5). We retrospectively used PhyDOSE at confidence levels *γ* ∈ {0.75, 0.95} to estimate the cells needed to perform an equivalent single-cell experiment. The mean of the per patient published false negative rate (*β* = 4.9%) was used to estimate the system error *a priori*. In the original study, a median of 7,584 cells per patient (IQR: 6,194-8,361) were sequenced. Fig 5e shows the distribution of PhyDOSE *k*\* for all patients with greater than one tree in the candidate set (median is 2, IQR: 2-6, max is 316) at *γ* ∈ {0.75, 0.95} versus the total number of cells sequenced in [34]. For *γ* = 0.95, the median value of *k*\* was 274 cells (IQR: 230-497). This is a significant reduction from the number of cells sequenced per patient in [34] with a median percent reduction at confidence level *γ* = 0.95 of 95.4% (IQR: 92.2%-98.0%) necessary to replicate the experiment with similar results (S1 Table).

### Prospective Analysis of a Non-small Cell Lung Cancer Cohort

Using PhyDOSE, we prospectively determined the number of cells needed to uniquely identify the true phylogeny for the 25 out of 100 patients in the TRACERx non-small-cell lung cancer cohort that have multiple candidate trees [3]. The authors previously identified the set 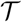 of candidate trees for each patient using CITUP [11] after clustering mutations with PyClone [36]. The authors also reported the cancer cell fraction of each mutation cluster in each bulk sample. The number of trees in the candidate set for each patient ranged from 2 to 17, with each containing mutation clusters with between 5 and 882 mutations (S2 Table).

Unlike in the simulations and ALL patient 2, multiple bulk samples per patient were available for analysis. Therefore, we calculated *k*\* for each sample independently for all 25 patients at varying confidence level *γ* ∈ {0.75, 0.95}. Mutation clusters alleviate the issue of false negatives, i.e. it suffices to only observe a single mutation to impute the presence of the other mutations in the same cluster. Here, with a typical SCS false negative rate of 0. 2, the probability of all mutations in the smallest cluster (with size 5) dropping out thus equals 0.2^5^ = 0.00032, a probability that can be neglected. As such, we set *β* = 0. The reported *k*\* value is the minimum *k*\* over the set of available samples, subsequently identifying which of the samples is the best to utilize for the SCS experiment. PhyDOSE was able to return a finite value of *k*\* for 23 out of the 25 patients.PhyDOSE will return ∞ when for each sample of the patient there is a featurette in every distinguishing feature where the clonal prevalance is 0. For two of the 23 patients the calculated *k*\* was over 400 due to featurettes in the distinguishing features with low clonal prevalence. For the remaining 21 patients, the median value of *k*\* was 29 for *γ* = 0.95 and 14 for *γ* = 0.75 (S9 Fig). These strikingly low values of *k*\* for the majority of the 25 patients with multiple candidate trees demonstrate the benefit of using PhyDOSE to strategically optimize the design of follow-up single cell experiments.

## Discussion

In this work, we showed that the mutation frequencies **f** and the set 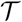 of tumor phylogenies inferred from initial bulk data contain valuable information to provide guidance for follow-up SCS experiments. We introduced PhyDOSE, a method to calculate the number *k*\* of single cells needed to infer the true phylogeny *T*\* given **f**, 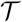 and a user-specified confidence level *γ*. Underpinning our method is the observation that often only a subset of clones suffices to distinguish one tree 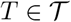 from the remaining trees 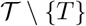.

We validated PhyDOSE using simulations and a retrospective analysis of leukemia patients [23, 34], concluding that PhyDOSE’s computed number *k*\* of cells resolves tree ambiguity, even in the presence of SCS errors. Our simulations showed that PhyDOSE remains robust in the presence of SCS errors such as elevated false negative and doublet rates as well as deviations between the distribution of clones in bulk and single-cell samples. Moreover, even in the case of an incomplete candidate tree set, PhyDOSE’s computed number *k*\* of cells sufficed to recover the true tree in a follow-up SCS experiment. In a prospective analysis, we demonstrated that only a small number of cells suffice to disambiguate the solution space of trees in a recent non-small cell lung cancer cohort [3]. Finally, we introduced the notion of support as a way to prioritize candidate trees given SCS data, often correctly selecting the ground truth tree among the candidate trees. Existing methods that infer tumor phylogeny from SCS data [14,16,32] or a combination of SCS and bulk data [20,33] may be used as an alternative. In summary, PhyDOSE proposes cost-efficient SCS experiments that will yield high-fidelity phylogenies, which may consequently improve downstream analyses in cancer genomics aimed at deepening our understanding of cancer biology.

There are several future research directions. First, in the case of multiple bulk samples, rather than selecting cells from a single sample, a better strategy would be to select cells across samples. To model this accurately, we must consider a multinomial mixture model. Second, to further reduce SCS costs, we might want to include a mutation selection step as part of our approach to perform targeted rather than whole-genome sequencing. Third, similar ideas can be used to design follow-up sequencing experiments using alternative sequencing technologies such as long read sequencing. Alternatively, performing additional bulk sequencing rather than single-cell sequencing might be more cost-effective, especially when obtaining a bulk sample with distinct clonal prevalences [10, 37]. Fourth, we plan to replace the integer linear program used for identifying minimal distinguishing features with a combinatorial algorithm. This will enable us to develop an easy-to-use and install R package with a Shiny user interface. Fifth, to improve robustness in the presence of SCS errors, we plan to explore alternative definitions of successful SCS experiment outcomes, requiring that more than one cells is observed of each featurette of a distinguishing feature. This will enable us to address errors such as doublets and false positives in a SCS experiment. Sixth, the concept of distinguishing features may be useful to summarize diverse solution spaces in cancer phylognetics [38]. Finally, we plan to explore evolutionary models beyond the infinite sites model, such as the Dollo parsimony model where mutations might be lost [16], requiring a more careful definition of the probability of observing a distinguishing feature.

## Supporting information

**S1 Fig. Reduction from Set Cover to *T*-SCS-PC**. Given a family 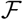 of subsets 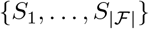 on a universe *U* = {1,…, *n*}, we construct *n* + 1 trees 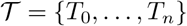 with mutations 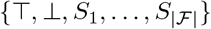 We seek to distinguish *T*_0_ from the remaining trees {*T*_1_, …, *T_n_*}. The key concept captured by the reduction is that there is a cover of size *k* if and only if Pr(*Y_k_* | **u**(*T*_0_, **f**)) is greater than 0. Here, *S*_1_ and *S*_4_ form a cover of size *k* = 2 of the universe U and the corresponding probability Pr(*Y_k_*| **u**(*T*_0_, **f**)) is greater than 0.

**S2 Fig. PhyDOSE sensitivity analysis of clonal prevalence distortion** (a) The mean absolute percentage difference of the single cell clonal prevalence per replication from simulated bulk clonal prevalence at values of λ ∈ {50, 2000}. These values of λ resulted in mean absolute percentage difference of 5% and 20% respectively. (b) λ = 2000 was selected for sim2a/b but further sensitivity analysis was performed with λ = 50. The recall metrics are compared between sim2a at λ = 2000 and sim2c at λ = 50. (b) Recall metrics when inferring *T*\* with SPhyR [16] by randomly sampling *k*\*/2, *k*\*, 2*k*\* simulated single cells.

**S3 Fig. Percentage of *in silico* single-cell sequencing experiments correctly identifying** *T*\* **as the true tree utilizing PhyDOSE’s** *k*\* **at** *γ* =0.95 and *β* ∈ {0.0, 0.2}. PhyDOSE has a high success rate in alignment with *γ* = 0.95 (dashed horizontal line) when utilizing the entire set 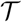 in condition *a* although the introduction of false negatives results in a reduction in success rate and greater variance. In the cases where 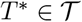, the support metric was still able to successfully identify *T*\* as the true tree.

**S4 Fig. Empirical runtime analysis on the bottleneck step of PhyDOSE** (a) Total runtime in seconds for each input set 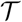 that completed in the two hour limit when solved sequentially. (b) Fraction of the input set 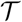 completed within the two hour limit when solved sequentially. (c) Mean runtime in seconds per tree 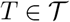.

**S6 Fig. Retrospective analysis of ALL patient 2 where [23] sequenced 115 single cells, targeting 16 mutations.** (a) Overview of single-cell data, focusing on a subset of 14 mutations. Each row is a single cell and columns correspond to mutations. Black cells indicate presence of the SNV, whereas white cells indicate absence. Text in each cell indicates a true positive (‘1’), false negative (‘1*’), true negative (‘0’) or false positive (‘0*’) entry. (b) Overview of the set 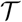 of trees, where each edge is labeled by its frequency in 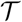. Red edges encode the tree *T*\* that is consistent with SCS data in (a). In particular, the SCS data encodes a tree composed of four mutation clusters, each indicated by a different shading.

**S7 Fig. Distribution of number of supporting cells for each candidate tree across 100 *in silico* SCS experiments at varying success probabilities for ALL patient 2 [23].** For *γ* = 0.75, we sampled *k*(*T*\*) = 50 cells from the SCS data. For *γ* = 0.95, we sampled *k*(*T*\*) = 103 cells. S8 Fig shows the candidate trees.

**S8 Fig. Candidate trees that had a non-zero support across 100 in *silico* SCS experiments for ALL patient 2 [23].** For *γ* = 0.75, we sampled *k*(*T*\*) = 50 cells from the SCS data. For *γ* = 0.95, we sampled *k*(*T*\*) = 103 cells. Labels match tree indices in S7 Fig.

**S1 Table Prospective analysis of TRACERx non-small-cell lung cancer cohort.** Table shows the patient identifier, the number of mutation clusters, the number of bulk samples, the minimum number of mutations per cluster, the maximum number of mutations per cluster, the size of the candidate set 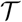 of trees as determined by [3], PhyDOSE’s *k*\* calculated at confidence levels of *γ* ∈ {0.75, 0.95} and the recommended sample label from which the single cells should be drawn.

**S2 Table Retrospective analysis of an acute myeloid leukemia (AML) cohort.** Table shows the patient identifier, the number of mutations, the total cells sequenced [34], the size of the candidate set 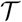 of trees as determined by SPRUCE [9], PhyDOSE’s *k*\* calculated at confidence levels of *γ* ∈ {0.75,0.95} (% reduction from total sequenced).

**S9 Fig. PhyDOSE calculated** *k*\* **for the lung cancer cohort at varying confidence levels.** Patients CRUK0013, CRUK0037, CRUK0068, CRUK0076 were excluded from the plot, but are shown in S1 Table.

**S1 Text Supplementary materials.**

## Acknowledgments

This work was supported by UIUC Center for Computational Biotechnology and Genomic Medicine (grant: CSN 1624790) and the National Science Foundation (grant: CCF 1850502).

## Supplementary Text

**Fig S1:**
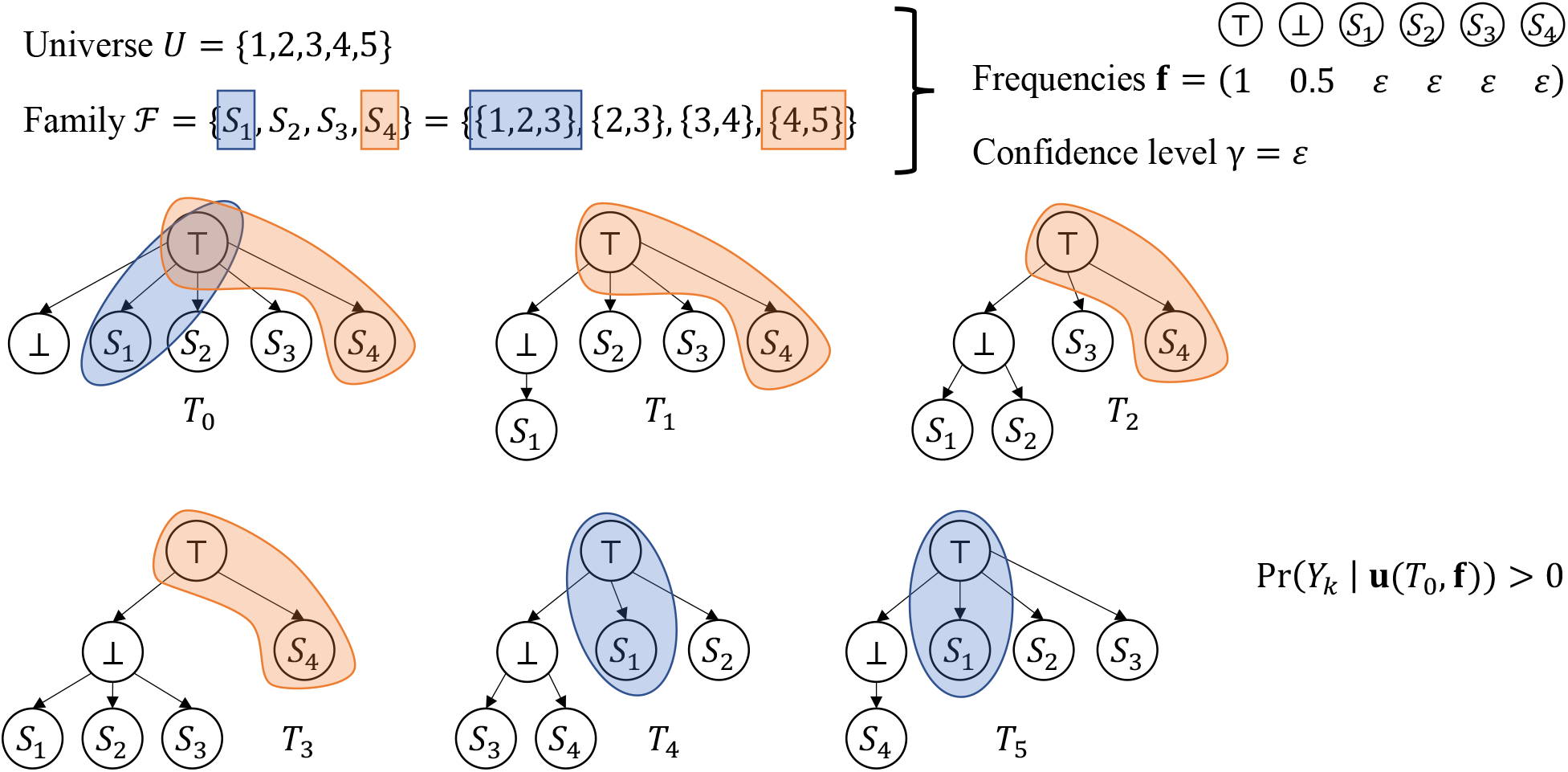
Reduction from Set Cover to *T*-SCS-PC. Given a family 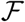 of subsets 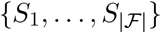 on a universe *U* = {1,…, *n*}, we construct *n* + 1 trees 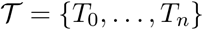 with mutations 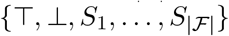 We seek to distinguish *T*_0_ from the remaining trees {*T*_1_,…, *T_n_*}. The key concept captured by the reduction is that there is a cover of size *k* if and only if Pr(*Y_k_*| **u**(*T*_0_, **f**)) is greater than 0. Here, *S*_1_ and *S*_4_ form a cover of size *k* = 2 of the universe *U* and the corresponding probability Pr(*Y_k_* | **u**(*T*_0_, **f**)) is greater than 0.

### A Supplementary Text

#### A.1 Complexity

##### Theorem 1.

*T*-SCS-PC is NP-hard.

We prove the theorem using a polynomial-time reduction from the Set Cover problem, a known NP-hard problem [1].

##### Problem 1

(Set Cover). Given a family 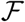 of subsets 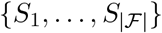 over a universe *U* = {1,…, *n*}, find a cover 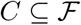 such that 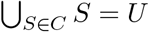 and *C* has minimum cardinality.

Specifically, we reduce a Set Cover instance 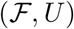 to an T-SCS-PC instance 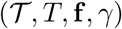 as follows. The set 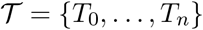 includes one tree *T_i_* for each element *i* in the universe *U* and an additional tree *T*_0_. All trees in 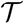 have 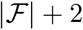 vertices, corresponding to subsets 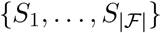 and two additional mutations {T, ⊥}. Each tree in 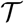 includes the edge (T, ⊥). Additionally, if element *i* ∈ *U* is absent from subset *S_j_* then there is an edge (T, *S_j_*) in tree *T_i_*, otherwise *T_i_* includes an edge (⊥, *S_j_*). Tree *T*_0_ includes edges (T, *S_j_*) for all subsets *S_j_*. As for the frequencies **f**, we set *f*_T_ = 1, *f*_⊥_ = 0.5 and the remaining frequencies *f_S_j__* = *ε* for all subsets 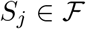. Moreover, we set the confidence level *γ* to *ε* as well. In the corresponding T-SCS-PC instance 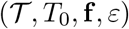, the tree of interest is *T*_0_. Fig. S1 shows an example.

The key idea is that as *γ* = *ε* > 0 is a small positive infinitesimal constant, this *T*-SCS-PC instance seeks the smallest number *k*\* of cells such that Pr(*Y_k*_* | **u**(*T*_o_, **f**)) is non-zero. In particular, this number *k*\* of cells will only be achieved if there is a distinguishing feature Π of the same size *k*\*. By our reduction, there is a 1-1 correspondence between set covers of *U* and distinguishing features Π of *T*_0_ with respect to {*T*_1_,…, *T_n_*}. Specifically, a set cover *C* of size *k* corresponds to a distinguishing feature Π(C) of the same size *k*, and vice versa. As such, we have the following lemma whose proof is in the supplement.

##### Lemma 1.

Let 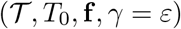 be the *T*-SCS-PC instance corresponding to Set Cover instance 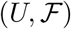. A minimum cover has size *k*\* if and only if *k*\* is the smallest integer such that Pr(*Y_k_*\* | **u**(*T*_0_, **f**)) ≥ *γ*.

*Proof. (⟹)* Let *C* be a minimum cover of the Set Cover instance 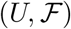. By the premise, we have that |*C*| = *k*\*. We start by showing that Pr(*Y_k_*\* | **u**(*T*_0_, **f**)) ≥ *γ* by constructing a distinguishing feature Π(*C*) of *T*_0_ where |Π| = *k*\*. Observe that for each subset *S_j_* in *C* we have that {T, *S_j_*} is a featurette of *T*_0_. We define Π(*C*) to be composed of featurettes {T, *S_j_*} for all subsets *S_j_* ∈ *C*. Thus, |⊓(*C*)| = *k*\*. To show that Π(*C*) is a distinguishing feature of *T*_0_, it remains to show that at least one featurette *τ* ∈ Π(*C*) is absent in each tree in 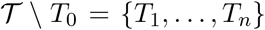. Consider any tree *T_i_* = *T*_0_. Since *C* is a cover, the element *i* of the universe *U* corresponding to tree *T_i_* must be covered by some subset *S_j_* ∈ *C*. This means that tree *T_i_* contains the edge (⊥, *S_j_*), which means that the featurette {T, *S_j_*} in Π(*C*) is absent from *T_i_*. Hence, Π(*C*) is a distinguishing feature of *T*_0_.

We now must show that Pr(*Y_k_*\* | **u**(*T*_0_, **f**)) ≥ *γ*. We do so by focusing on distinguishing feature Π(*C*). By construction of *T*_0_ and **f**, it follows from (1) that each featurette {T, *S_j_*} in Π(*C*) has a clonal prevalence *U_j_* = *ε*. This means that a SCS experiment of *k*\* cells where we only observe the *k*\* featurettes/clones has a probability that is strictly greater than 0. Therefore, Pr(*Y_k_*\* | **u**(*T*_0_, **f**)) > 0. Since *ε* is a small positive infinitesimal constant, we have that Pr(*Y_k*_* | **u**(*T*_0_, **f**)) ≥ *γ* = *ε*.

It remains to show that *k*\* is the smallest integer where Pr(*Y_k*_* | **u**(*T*_0_, **f**)) ≥ *ε*. Assume for a contradiction that the smallest integer *k*’ where Pr(*Y_k_* | **u**(*T*_0_, **f**)) ≥ *ε* is strictly smaller than *k*\*. This means that there exists a minimal distinguishing feature Π’ of size at most k’. By definition Π’ is composed of featurettes corresponding to root-to-vertex paths in *T*_0_. Since Π’ is minimal, it will not contain the featurette {T, ⊥} as this featurette is present in all remaining trees {*T*_1_,…, *T_n_*}. Thus, Π’ is composed of featurettes of the form {T, *S_j_*} where 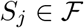. Since Π’ is a distinguishing feature, no tree *T*_1_ ∈ {*T*_1_,…, *T_n_*} contains all featurettes of Π’. By construction of {*T*_1_,…, *T_n_*}, this means that the subsets encoded in Π’ form a cover of the universe *U*. Thus, there exists a cover with size strictly smaller than *k*\*, contradicting the premise. Therefore, *k*\* is indeed the smallest integer where Pr(*Y_k*_* | **u**(*T*_0_, **f**)) ≥ *γ = ε*.

(⇐) Let *k*\* be the smallest integer such that Pr(*Y_k*_* | **u**(*T*_0_, **f**)) ≥ *γ* = *ε*. We start by showing that the size of a minimum distinguishing feature Π of *T*_0_ has to be exactly *k*\*. Clearly, if |Π| > *k*\* then Pr(*Y_k*_* | **u**(*T*_0_, **f**)) = 0 as there exists no successful SCS experiment with *k*\* cells. On the other hand, if |Π| < *k*\* then there exists a successful SCS experiment with |Π| cells. In other words, Pr(*Y*_|Π|_ |**u**(*T*_0_, **f**)) ≥ *ε*. This contradicts that *k*\* is the smallest integer where Pr(*Y*_|Π|_ | **u**(*T*_0_, **f**)) ≥ ε. Hence, |Π| = *k*\*.

Consider a minimum distinguishing feature Π of *T*_0_. By the previous argument, we know that |Π| = *k*\*. We will show that Π encodes a cover *C*(Π) of *U* of size *k*\*. Since Π is minimal, it will not contain the featurette {T, ⊥} of *T*_0_ as this featurette is present in all remaining trees {*T*_1_,…, *T_n_*}. Thus, Π is composed of *k* featurettes of the form {T, *S_j_*} where 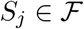. Let *C*(Π) be defined as the collection of subsets 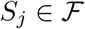 where {T, *S_j_*} in Π. Since Π is a distinguishing feature, no tree *T_i_* ∈ {*T*_1_,…, *T_n_*} contains all featurettes of Π. By construction of {*T*_1_,…, *T_n_*}, this means that *C*(Π) is a cover of size *k* of the universe *U*.

Finally, we must show that there exists no cover *C*’ of *U* with size |*C*’| strictly smaller than *k*\*. Suppose for a contradiction that such a cover *C*’ exists. By construction, *C*’ encodes a distinguishing feature Π(*C*’) composed of featurettes {T, *S_j_*} for all subsets *S_j_* ∈ *C*’. Thus, |Π(*C*’) | = |*C*’|. To show that Π(*C*’) is a distinguishing feature of *T*_0_, we must show that (i) all features *τ* ∈ Π(*C*’) are present in *T*_0_, and (ii) at least one featurette *τ* ∈ Π(*C*) is absent in each tree in 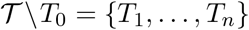. Condition (i) holds by construction of Π(*C*’) and *T*_0_, i.e. for each subset *S_j_* in *C*’ we have that {T, *S_j_*} is a featurette of *T*_0_. As for condition (ii), consider any tree *T_i_* = *T*_0_. Since *C*’ is a cover, the element *i* of the universe *U* corresponding to tree *T_i_* must be covered by some subset *S_j_* ∈ *C*’. This means that tree *T_i_* contains the edge (⊥, *S_j_*), which means that the featurette {T, *S_j_*} in Π(*C*’) is absent from *T_i_*. Hence, Π(*C*’) is a distinguishing feature of *T*_0_. This in turn means that Pr(*Y*_|Σ|(*C*’)_ | **u**(*T*_0_, **f**)) > 0. In other words, Pr(*Y*_|Σ|(*C*’)_ | **u**(*T*_0_, **f**)) ≥ *γ* = *ε*, thus contradicting the premise. Hence, minimum set covers of 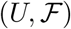 have cardinality *k*\*.

The theorem follows from the above lemma, as the reduction to obtain 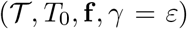 from 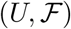 takes only polynomial time.

### B Supplementary Methods

#### B.1 Finding the Minimal Distinguishing Feature Family Φ*

To perform the calculation in (4), it is necessary to first find the minimal distinguishing feature family Φ*. Using similar ideas as in our hardness proof (Section A.1), we consider the reverse reduction from the problem of finding a minimal distinguishing feature to that of finding a minimum size set cover (Problem 1).

We define the universe *U* = {1,…,*m*} to be the set 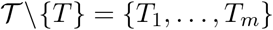 of trees excluding the tree *T* for which we want to solve the *T*-SCS-PC problem. We define the family 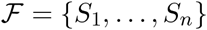 of subsets to correspond to the *n* featurettes present in *T*. Specifically, the subset *S_j_* corresponding to featurette *τ_j_* that is present in *T* is composed of elements *i ∈ U* corresponding to trees *T_i_* where *τ_j_* is absent. The key idea is that *S_j_* is indicating in which input trees featurette *τ_j_* of *T* is absent. We note that 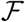 is a multi-set as distinct featurettes *τ_j_* and *τ_j’_* may be absent in the same set of trees, thus leading to *S_j_* = *S_j’_* (Fig. 3(b)). There is a bijection between set covers of 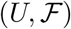 and distinguishing features of *T*. That is, each distinguishing feature Π = {*τ*_1_,…, *τ*_|π|_} corresponds to the same-sized cover Π(*C*) composed of subsets {*S*_1_,…, *S*_|Π|(*C*)_ }, and vice versa. In particular, a minimal distinguishing feature corresponds to a minimal set cover. Thus, we may use the following integer linear program (ILP) to find a minimum set cover *C* and thus a corresponding *minimum* distinguishing feature Π(*C*) — a minimal distinguishing feature of minimum size.

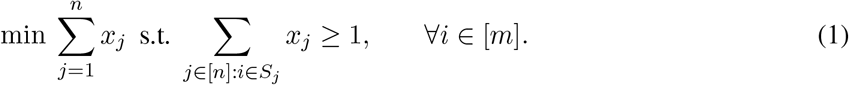

In order to find the *next* minimum distinguishing feature Π(*C*’) that is not contained within Π(*C*), we add the following constraint to the ILP.

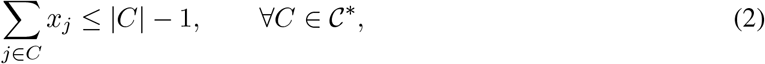

where 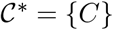. By repeatedly adding identified minimum set covers to 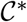 until the ILP becomes infeasible, we identify all minimal set covers 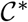 and thus all minimal distinguishing features Φ*. Fig. 3(b) shows an example. We use IBM ILOG CPLEX v12.9 to solve the ILP^1^.

### C Supplementary Results

**Fig S2:**
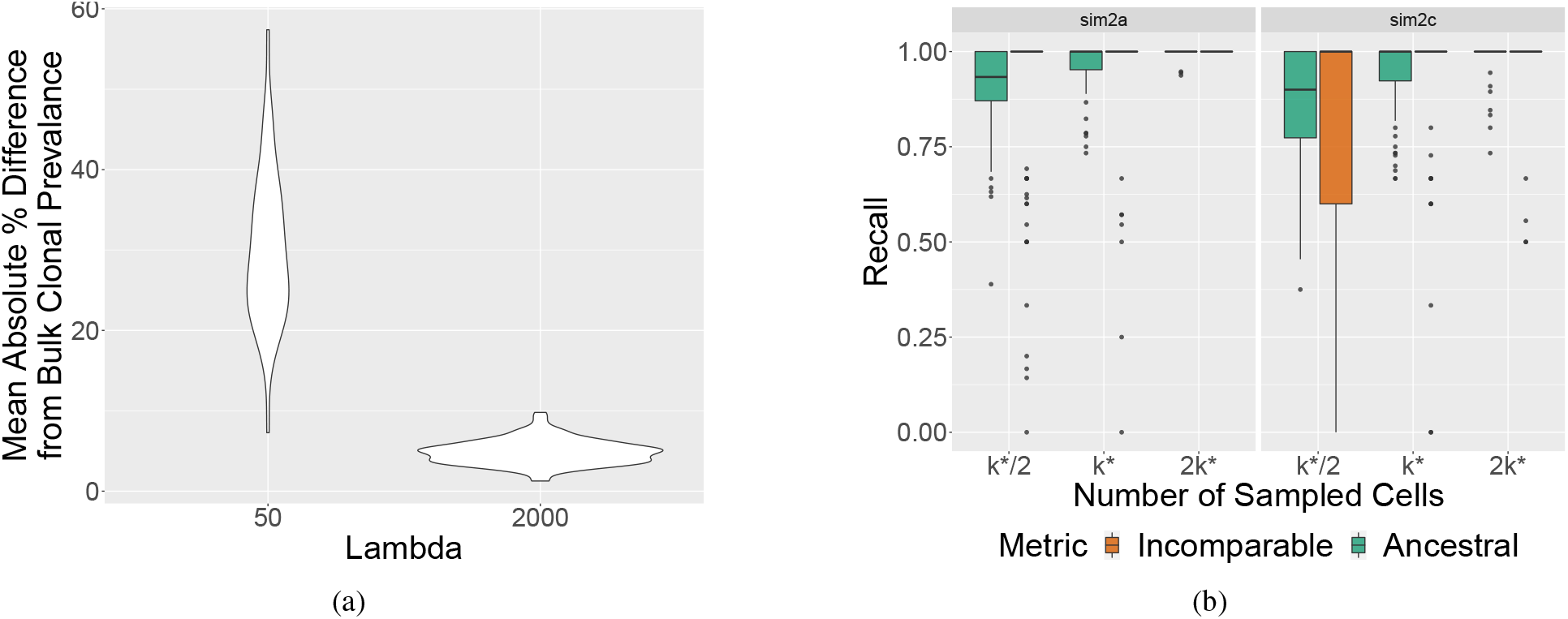
PhyDOSE sensitivity analysis of clonal prevalence distortion. (a) The mean absolute percentage difference of the single cell clonal prevalence per replication from simulated bulk clonal prevalence at values of λ ∈ {50,2000}. These values of λ resulted in mean absolute percentage difference of 5% and 20% respectively. (b) λ = 2000 was selected for sim2a/b but further sensitivity analysis was performed with λ = 50. The recall metrics are compared between sim2a at λ = 2000 and sim2c at λ = 50. (b) Recall metrics when inferring *T*\* with SPhyR [2] by randomly sampling *k*\*/2, *k*\*, 2*k*\* simulated single cells.

**Fig S3:**
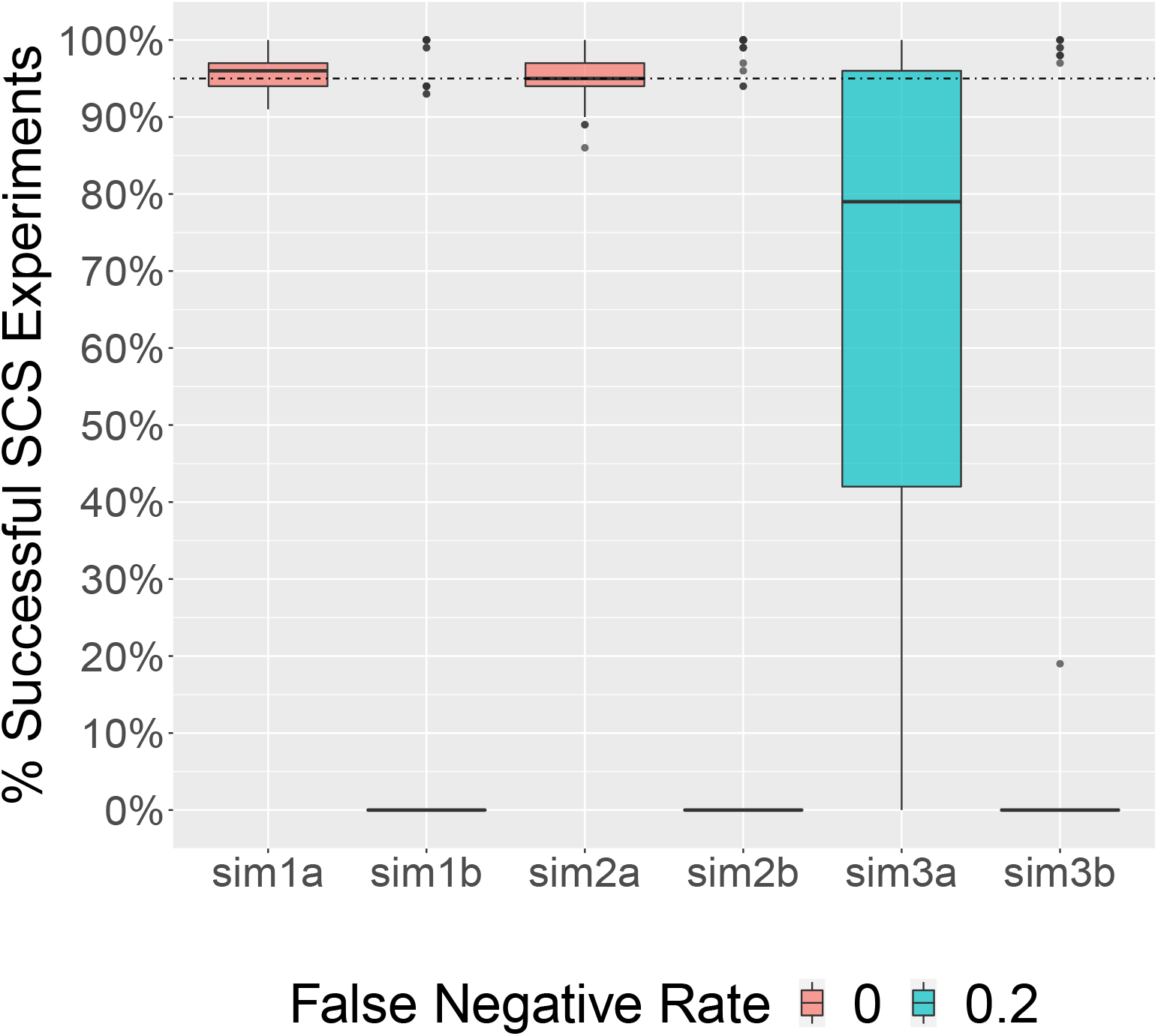
Percentage of *in silico* single-cell sequencing experiments correctly identifying T* as the true tree utilizing PhyDOSE’s *k*\* at γ =0.95 and *β* ∈ {0.0,0.2}. PhyDOSE has a high success rate in alignment with *γ* = 0.95 (dashed horizontal line) when utilizing the entire set 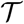 in condition *a*. However, the introduction of false negatives results in a reduction in success rate and greater variance. In the cases where 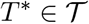, the support metric was still able to successfully identify *T*\* as the true tree.

**Fig S4:**
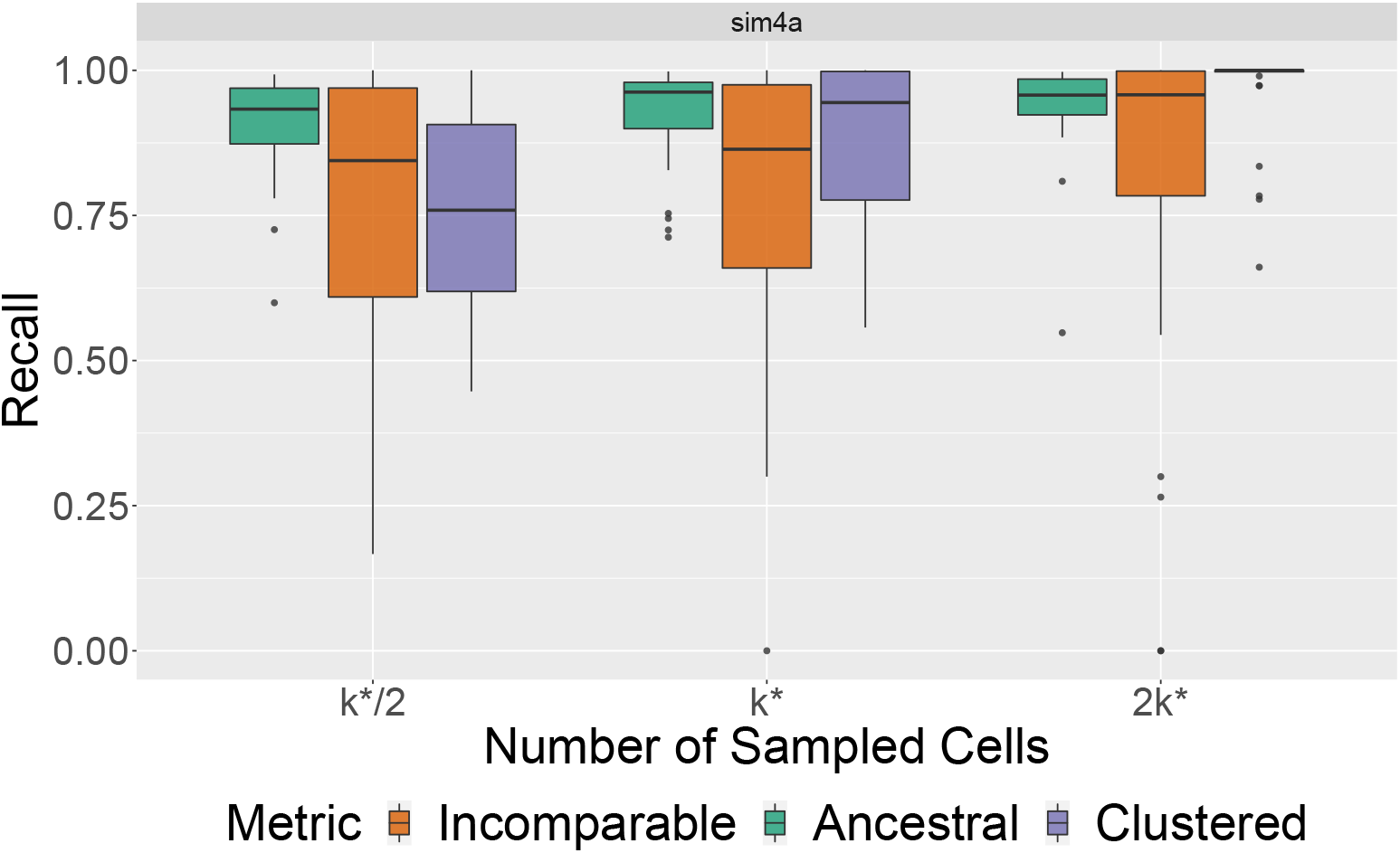
PhyDOSE performance with mutation clustering. Recall metrics when inferring *T*\* with SPhyR [2] by randomly sampling *k*\*/2, *k*\*, 2*k*\* simulated single cells.

**Fig S5:**
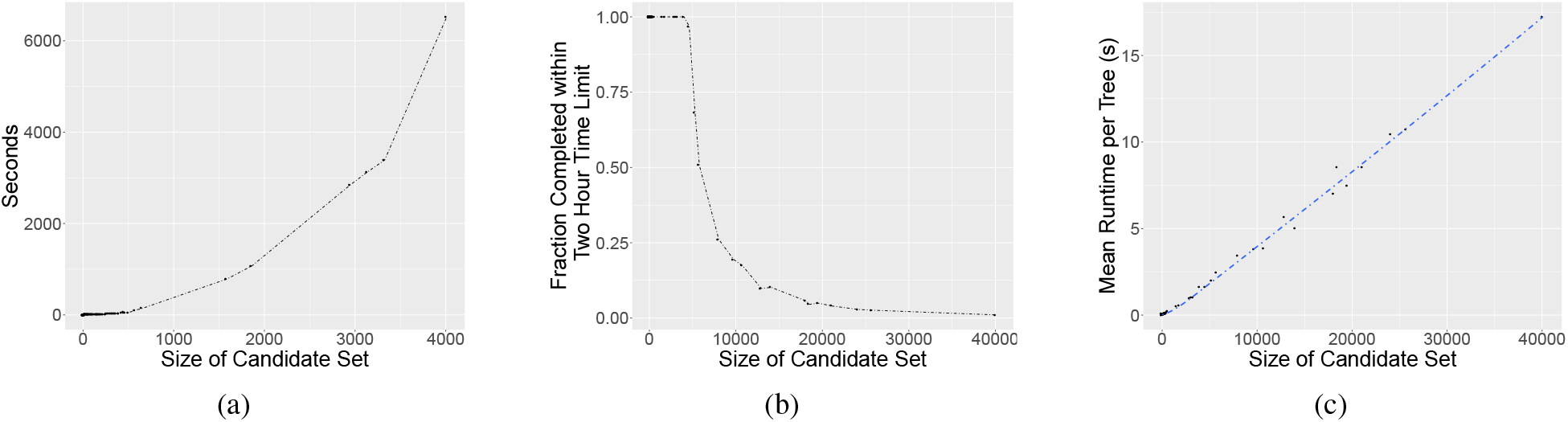
Empirical runtime analysis on the bottleneck step of PhyDOSE. (a) Total runtime in seconds for each input set 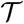 that completed in the two hour limit when solved sequentially. (b) Fraction of the input set 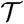 completed within the two hour limit when solved sequentially. (c) Mean runtime in seconds per tree 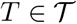.

#### C.1 Retrospective Analysis of an Acute Lymphoblastic Leukemia Patient

For each patient, the sequenced cells are clustered into 2 to 7 clones using an EM-based approach [3]. Based on the fact that false negatives occur more frequently than false positives, we designated an SNV as present if at least 30% of cells in the clone had the mutation. We then checked if the resulting binary, clone-by-SNV matrices adhered to the infinite sites assumption, which was the case for only patients 2 and 3. While the VAFs of all 16 SNVs in patient 2 are less than 0.5, patient 3 had 6 out 49 SNVs with a VAF larger than 0.5, which is indicative of copy number aberrations. Since no copy number information was available to infer cancer cell fractions, we excluded patient 3 from our analysis, thus restricting our attention to patient 2.

For patient 2, 115 cells were clustered into 5 clones [3]. This patient has 16 SNVs, from which we excluded mutations *CMTM8, ATRNL1, L1NC00052* and *TRRAP* for reasons that we described in the main text. The majority voting rule described above yielded a binary clone-by-SNV matrix with 4 mutations clusters that each correspond to SNVs that co-occur in every clone (Fig. S6a), corresponding to a two-state perfect phylogeny *T*_SES_ on the mutation clusters (Fig. 5(b)). To obtain the set 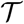 of candidate phylogenies, we considered the bulk data. Specifically, we merged mutations ZC3H3 and XPO7 as they had the same VAF in the bulk data and occurred in the same mutation cluster in the cleaned SCS data (Fig. 5(a)). Using SPRUCE [4], we enumerated 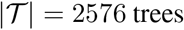 trees (Fig. S6b). Only one tree 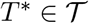 was consistent with *T*_SES_, i.e. each mutation cluster of *T*_SES_ formed a connected path in *T*\* and subsequently collapsing these paths in *T*\* yields *T*_SES_. Comparing the cleaned single-cell data to the raw values, we computed a false negative rate *β* of 0.2 for the 14 mutations (Fig. S6a), which was in line with the value reported by [3].

**Fig S6:**
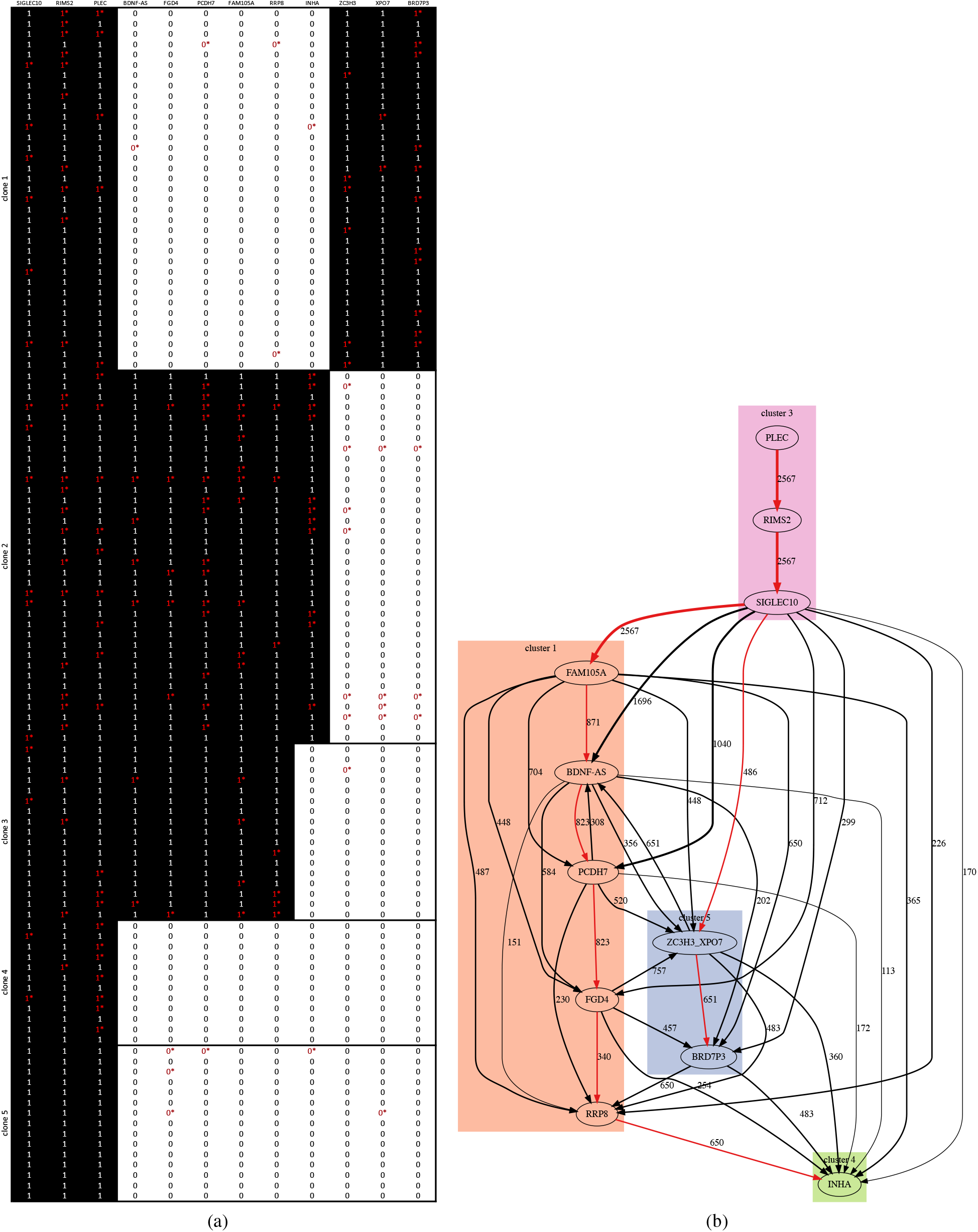
Retrospective analysis of ALL patient 2 where [3] sequenced 115 single cells, targeting 16 mutations. (a) Overview of single-cell data, focusing on a subset of 14 mutations. Each row is a single cell and columns correspond to mutations. Black cells indicate presence of the SNv, whereas white cells indicate absence. Text in each cell indicates a true positive (‘1’), false negative (‘1*’), true negative (‘0’) or false positive (‘0*’) entry. (b) Overview of the set 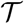 of trees, where each edge is labeled by its frequency in 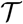. Red edges encode the tree *T*\* that is consistent with SCS data in (a). In particular, the SCS data encodes a tree composed of four mutation clusters, each indicated by a different shading.

**Fig S7:**
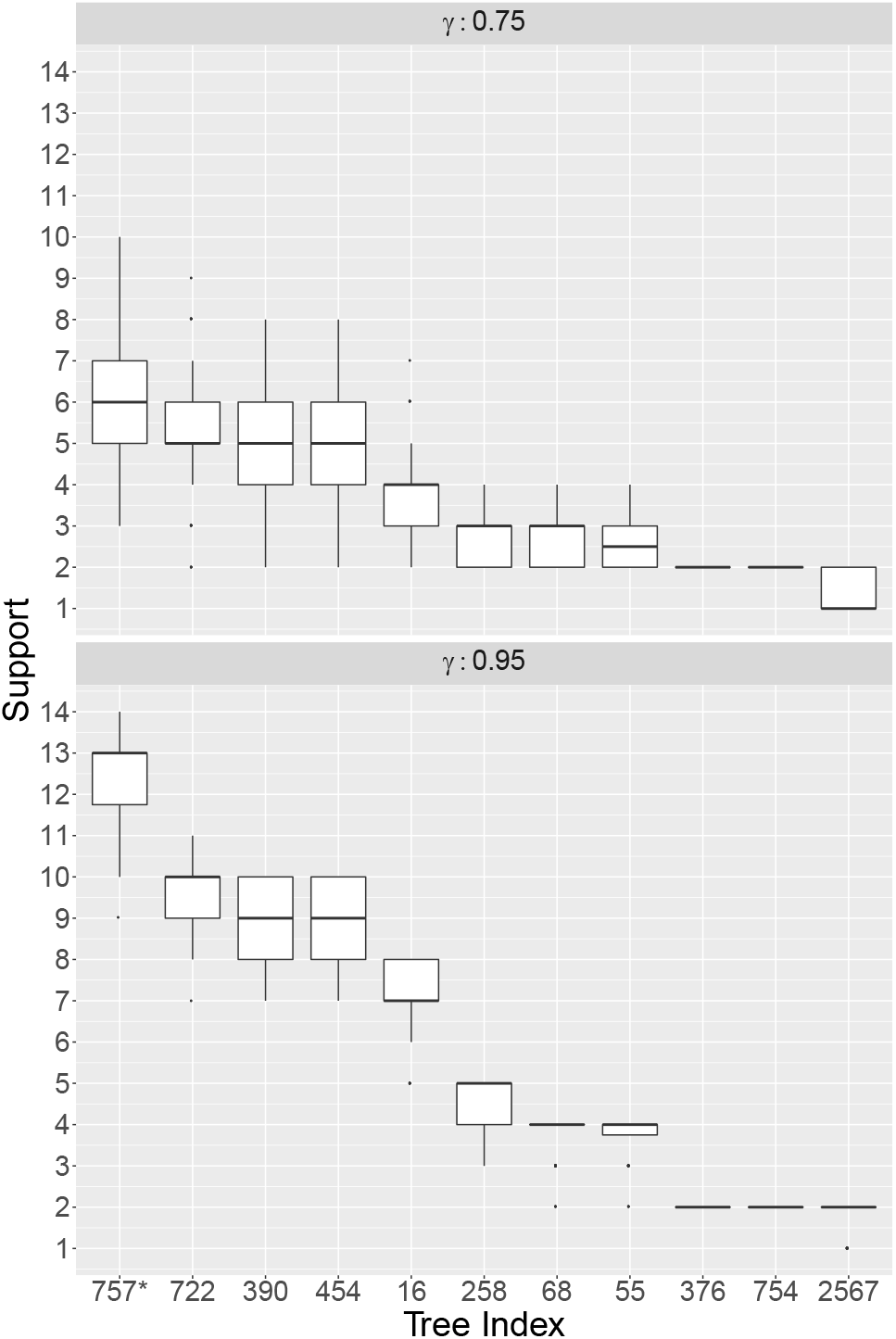
Distribution of number of supporting cells for each candidate tree across 100 in *silico* SCS experiments at varying success probabilities for ALL patient 2 [3]. For *γ* = 0.75, we sampled *k*(*T*\*) = 50 cells from the SCS data. For *γ* = 0.95, we sampled *k*(*T*\*) = 103 cells. Fig. S8 shows the candidate trees.

**Fig S8:**
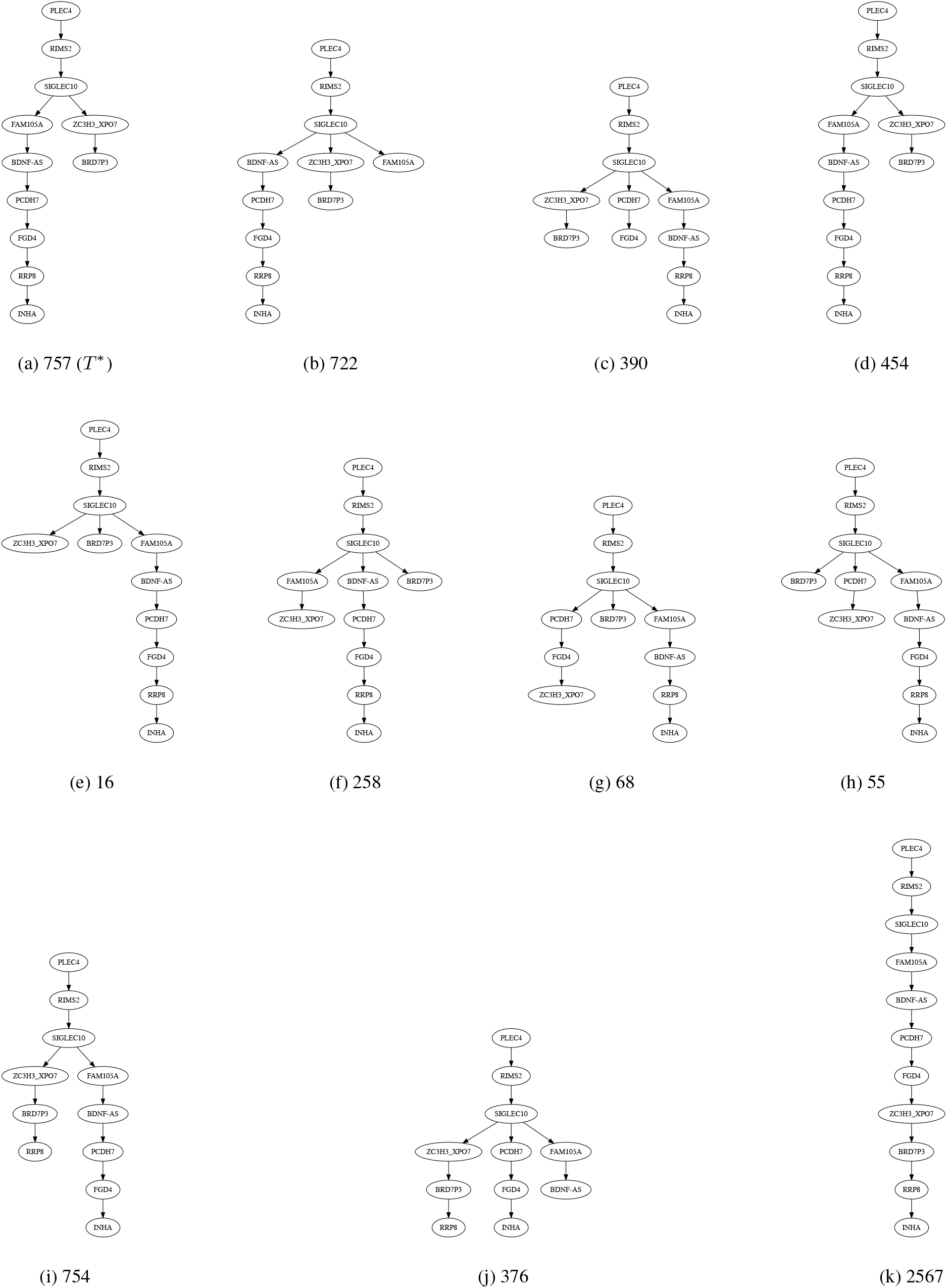
Candidate trees that had a non-zero support across 100 in *silico* SCS experiments for ALL patient 2 [3]. For *γ* = 0.75, we sampled *k*(*T*\*) = 50 cells from the SCS data. For *γ* = 0.95, we sampled *k*(*T*\*) = 103 cells. Labels match tree indices in Fig. S7.

**Table S1:**
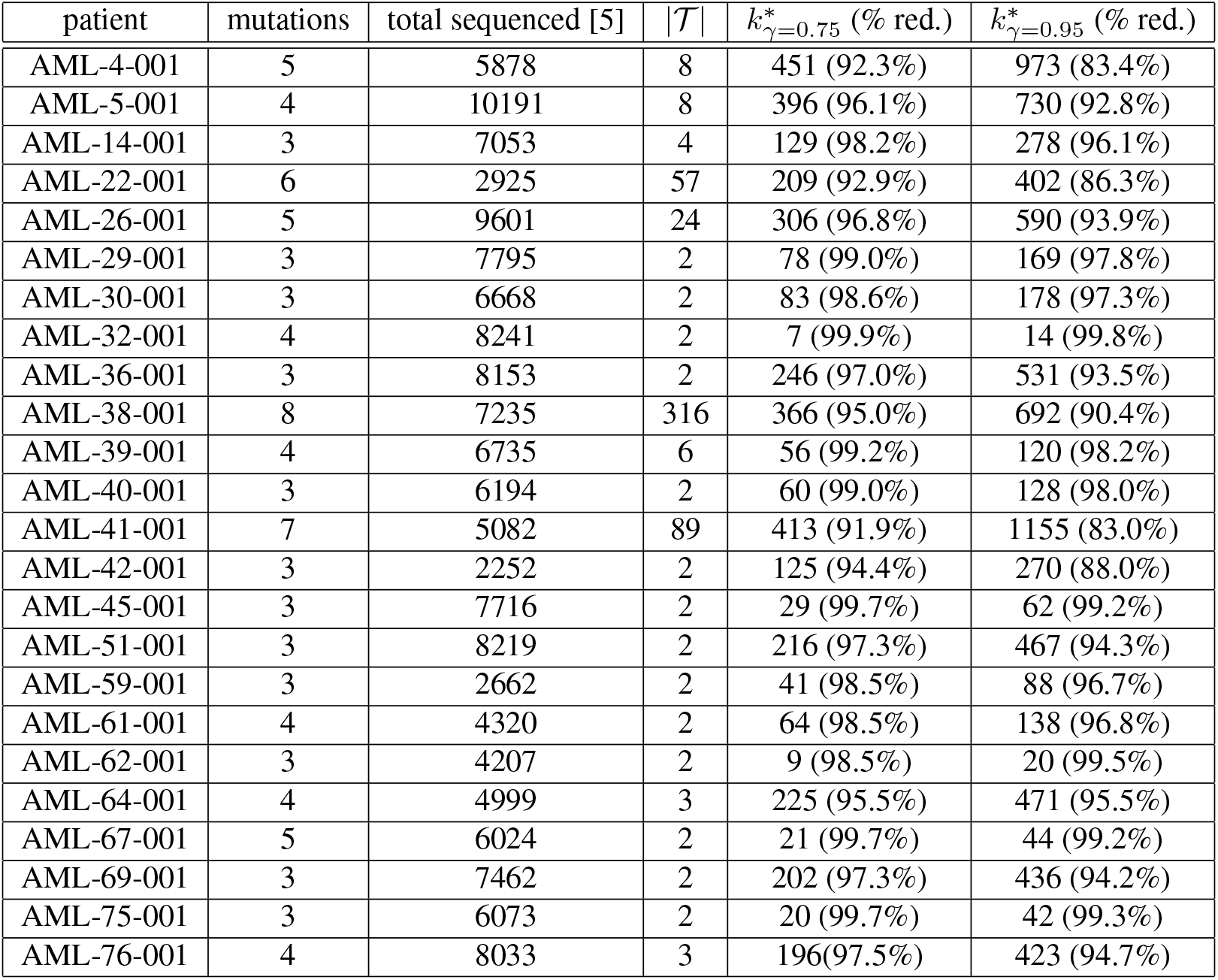
Prospective analysis of an acute myeloid leukemia (AML) cohort. Table shows the patient identifier, the number of mutations,the total cells sequenced [5], the size of the candidate set 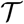 of trees as determined by SPRUCE [4], PhyDOSE’s *k*\* calculated at confidence levels of *γ* ∈ {0.75,0.95} (% reduction from total sequenced).

**Table S2:** Prospective analysis of TRACERx non-small-cell lung cancer cohort. Table shows the patient identifier, the number of mutation clusters, the number of bulk samples, the minimum number of mutations per cluster, the maximum number of mutations per cluster, the size of the candidate set 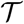 of trees as determined by [6], PhyDOSE’s *k*\* calculated at confidence levels of *γ* ∈ {0.75, 0.95} and the recommended sample label from which the single cells should be drawn.

| patient | clusters | samples | min muts | max muts | $\vert \mathcal{T}\vert $ | $k^*(\gamma = 0.75)$ | $k^*(\gamma = 0.95)$ | sel. sample |
| --- | --- | --- | --- | --- | --- | --- | --- | --- |
| CRUK0004 | 7 | 4 | 10 | 78 | 2 | 15 | 32 | R3 |
| CRUK0005 | 6 | 4 | 24 | 536 | 2 | 12 | 26 | R3 |
| CRUK0011 | 8 | 3 | 12 | 335 | 3 | 16 | 34 | R2 |
| CRUK0012 | 5 | 2 | 11 | 84 | 2 | 7 | 15 | R1 |
| CRUK0013 | 9 | 5 | 5 | 114 | 8 | 487 | 1051 | LN1 |
| CRUK0022 | 5 | 2 | 5 | 158 | 2 | 11 | 23 | R1 |
| CRUK0023 | 10 | 4 | 5 | 226 | 2 | 10 | 20 | R1 |
| CRUK0025 | 7 | 3 | 10 | 350 | 2 | 12 | 25 | R2 |
| CRUK0028 | 5 | 2 | 5 | 72 | 2 | 15 | 33 | R1 |
| CRUK0031 | 7 | 3 | 15 | 675 | 2 | 14 | 30 | R1 |
| CRUK0037 | 10 | 5 | 6 | 397 | 17 | ∞ | ∞ | N/A |
| CRUK0038 | 4 | 2 | 6 | 107 | 2 | 20 | 43 | R1 |
| CRUK0046 | 5 | 4 | 5 | 186 | 2 | 14 | 29 | R1 |
| CRUK0049 | 6 | 2 | 7 | 882 | 4 | 17 | 37 | R2 |
| CRUK0063 | 8 | 5 | 5 | 167 | 2 | 16 | 33 | R4 |
| CRUK0067 | 5 | 2 | 24 | 263 | 2 | 14 | 29 | R1 |
| CRUK0068 | 10 | 4 | 11 | 532 | 3 | ∞ | ∞ | N/A |
| CRUK0070 | 10 | 5 | 6 | 254 | 2 | 11 | 24 | R6 |
| CRUK0076 | 9 | 4 | 8 | 846 | 4 | 21971 | 47470 | R2 |
| CRUK0077 | 7 | 4 | 9 | 586 | 2 | 10 | 22 | R4 |
| CRUK0084 | 6 | 4 | 5 | 332 | 2 | 8 | 17 | R2 |
| CRUK0094 | 6 | 4 | 6 | 50 | 2 | 10 | 22 | R4 |
| CRUK0095 | 4 | 3 | 6 | 216 | 2 | 20 | 42 | R2 |
| CRUK0099 | 5 | 4 | 5 | 438 | 2 | 21 | 46 | R6 |
| CRUK0100 | 8 | 3 | 5 | 777 | 3 | 12 | 24 | R2 |

**Fig S9:**
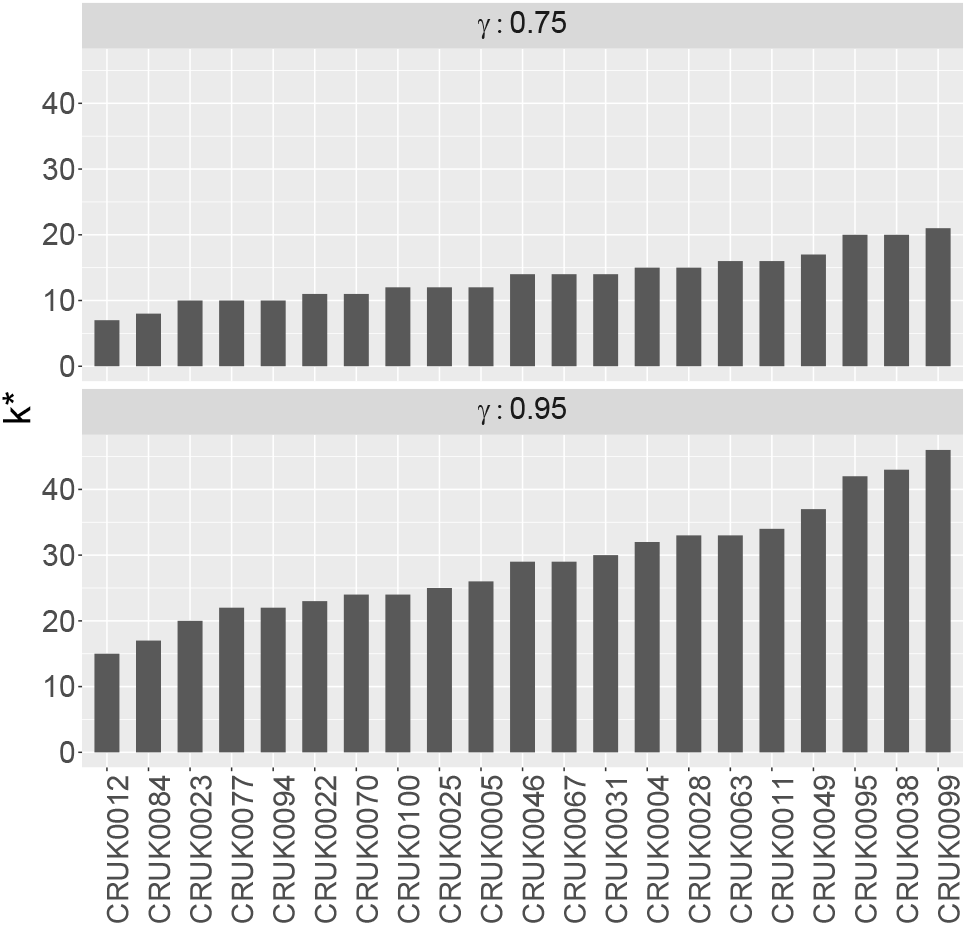
PhyDOSE calculated *k*\* for the lung cancer cohort at varying confidence levels. Patients CRUK0013, CRUK0037, CRUK0068, CRUK0076 were excluded from the plot, but are shown in Table S2.

## Footnotes

1 https://www.ibm.com/analytics/cplex-optimizer

